# Self-quenched fluorophore-DNA labels for super-resolution fluorescence microscopy

**DOI:** 10.1101/2024.03.24.586443

**Authors:** Laurell Kessler, Ashwin Balakrishnan, Tanja Menche, Dongni Wang, Yunqing Li, Maximilian Mantel, Marius Glogger, Marina S. Dietz, Mike Heilemann

**Author notes:** contributed equally (Laurell Kessler, Ashwin Balakrishnan).

## Abstract

Protein labeling through transient and repetitive hybridization of short, fluorophore-labeled DNA oligonucleotides has become widely applied in various optical super-resolution microscopy methods. The main advantages are multi-target imaging and molecular quantification. A challenge is the high background signal originating from the presence of unbound fluorophore-DNA labels in solution. Here, we report self-quenching of fluorophore dimers conjugated to DNA oligonucleotides as a general concept to reduce the fluorescence background. Upon hybridization, the fluorescence signal of both fluorophores is fully restored. Here, we expand the toolbox of fluorophores suitable for self-quenching and report their spectra and hybridization equilibria. We apply self-quenched fluorophore-DNA labels to stimulated emission depletion (STED) microscopy and single-molecule localization microscopy (SMLM) and report improved imaging performances.

## Introduction

Fluorophore-labeled, short DNA oligonucleotides are extensively used in various super-resolution microscopy applications, including the single-molecule localization microscopy (SMLM) ^1^ method DNA points accumulation in nanoscale topography (DNA-PAINT) ^2^, super-resolution optical fluctuation imaging (SOFI) ^3^ and stimulated emission depletion (STED) microscopy ^4,5^. An advantage that applies to all these applications is multi-target imaging, which is enabled by repetitive rounds of imaging and washing of sequence-orthogonal DNA strands. An additional benefit that is particular to single-molecule DNA-PAINT is the opportunity for molecular quantification by kinetic analysis ^6^. Furthermore, weak-affinity DNA labels were shown to optimize the performance of neural networks that predict super-resolved images from high-density single-molecule data ^7^.

Despite all these opportunities, the use of weak-affinity DNA labels demands keeping them in the imaging buffer during a microscopy experiment, which results in a higher fluorescence background and a limit in acquisition speed. To address these challenges, FRET-quenched fluorophores attached to extended DNA oligonucleotides were introduced ^8^. Recently, DNA oligonucleotides labeled with two identical oxazine fluorophores on either end were shown to exhibit self-quenching through short-distance molecular interactions ^9,10^. These probes led to a lower background signal in the unbound state, and consequently to a higher signal-to-background (SBR) ratio. In addition, an almost two-fold higher photon yield in the bound state increased the localization precision (DNA-PAINT) and spatial resolution (DNA-PAINT and STED-PAINT) ^9^. Here, we generalize this approach and introduce new fluorophores to exploit self-quenching with short DNA oligonucleotides. We characterize the spectroscopic and hybridization properties, demonstrate STED and DNA-PAINT imaging, and discuss a strategy for an experiment-driven general design.

## Results

We designed fluorophore-labeled DNA oligonucleotides (“imager strands”) using a well-established sequence frequently used for DNA-PAINT (P1; see **Table S1**) ^2^ and carrying either two fluorophores at the 5’- and 3’-end (dye-P1-dye), or one fluorophore at the 3’-end (P1-dye) (**Figure 1A**). As fluorophores, we selected one representative out of three classes of organic fluorophores that are used in advanced fluorescence microscopy methods: the carbocyanine Cy3B, a silicon-rhodamine (SiR) and tetramethylrhodamine (TMR) (**Table S1**). We characterized these fluorophore-DNA probes as single-stranded oligonucleotides, as well as hybridized to a sequence-complementary DNA oligonucleotide (“docking strand”), using absorption and fluorescence spectroscopy (**Figure 1B**). The absorption spectra of dual-labeled, single-stranded Cy3B-P1-Cy3B and TMR-P1-TMR showed a clear blue-shifted band, which indicates the formation of a fluorophore dimer; this band disappeared when these probes were hybridized to the docking strand (**Figure 1Bi**). For SiR-P1-SiR, only a faint dimer band was detected (**Figure 1Bi**). The fluorescence emission spectra of all three dual-labeled, single-stranded probes showed an effective increase in fluorescence upon hybridization of 3.3 (SiR-P1-SiR), 5.2 (TMR-P1-TMR), and 1.7 (Cy3B-P1-Cy3B) (**Figure 1Bii**). This indicates quenching of dual-labeled imager strands when freely diffusing in solution. In order to assess the kinetics of fluorescence quenching, we recorded fluorescence correlation spectroscopy (FCS) data of single- and dual-labeled probes and observed multiple sub-diffusional kinetics (**Figure S1, Table S2**). One of the sub-diffusional kinetics, in the range of 10^-7^ s, was only observed for dual-labeled imager strands and was absent for single-labeled counterparts (**Figure S2**). For the dual-labeled probes, this fast kinetic component appears only for freely diffusing imager strands and completely disappears when hybridized to docking strands hinting that it arises from the self-quenching process (**Figure S2, red**). We further quantified the k_fluorescence_ and k_dark_ rates of the self-quenching process in the range of 10^5^-10^6^ s^-1^ (**Figure S3, Table S2**).

**Figure 1.**
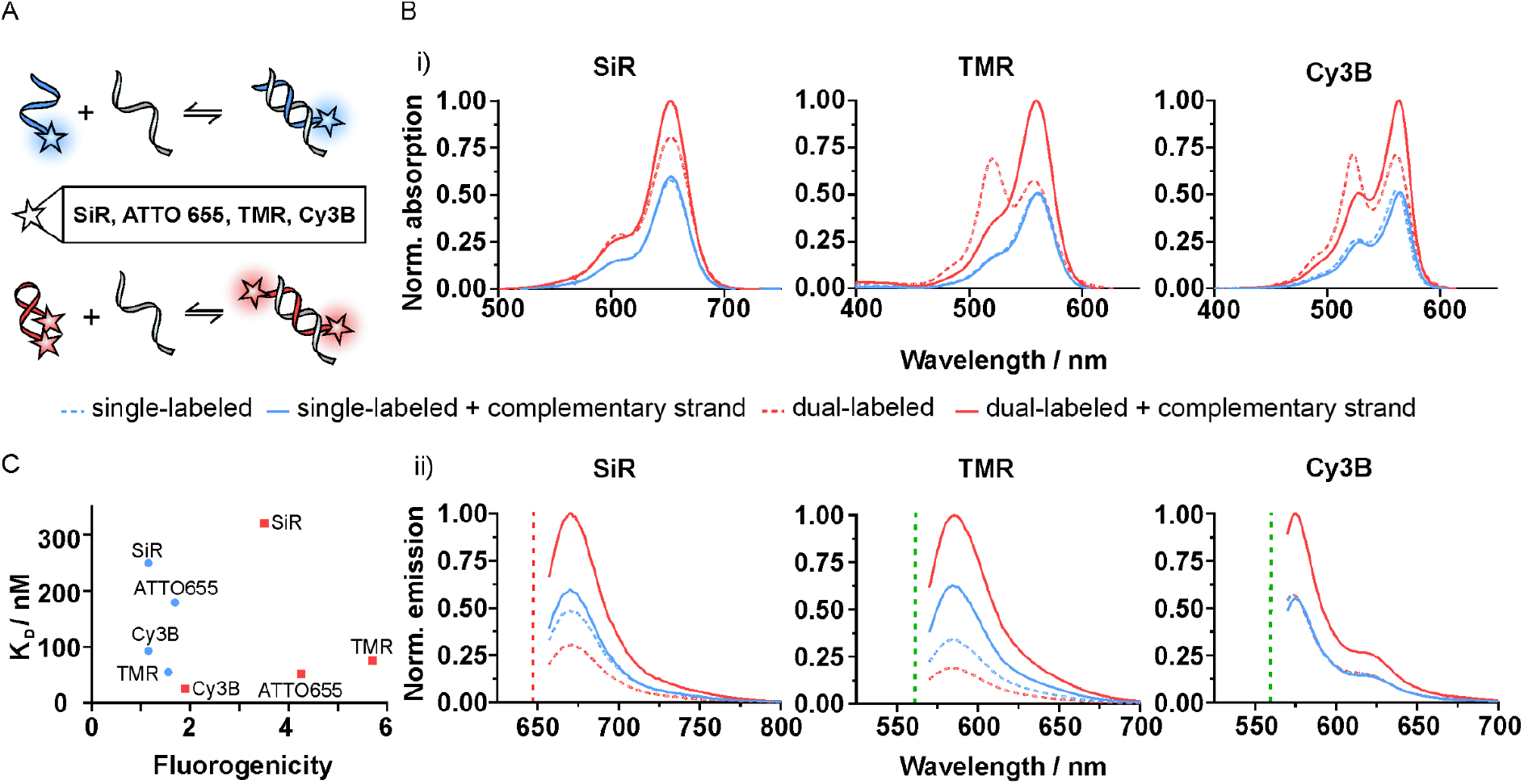
Spectroscopic characterization and hybridization equilibria for single- and dual-labeled DNA oligonucleotides. **A**. Schematic of conventional DNA-PAINT (blue) and DNA-PAINT using self-quenching fluorophores (red). Inlay shows the fluorophores that exhibit self-quenching. **B**. (i) Absorption and (ii) emission spectra of single-(3’) and dual-labeled (3’ and 5’) P1 imager strand sequence labeled with SiR, TMR, and Cy3B diluted in DNA-PAINT imaging buffer with and without sequence-complementary docking strands. Excitation lasers used for emission spectra measurement are shown with dashed lines in B, red: 647 nm; green: 560 nm. **C**. Fluorogenicity plotted against K_D_ of dual-labeled (red) and single-labeled (blue) P1 imager strands of different fluorophores used in this study and that of ATTO 655.

Next, we compared the hybridization properties of single- and dual-labeled DNA oligonucleotides by measuring the dissociation constant (K_D_). We found an increased binding affinity (decreased K_D_) in the case of Cy3B-P1-Cy3B relative to P1-Cy3B, and a slightly decreased affinity in the case of both SiR-P1-SiR and TMR-P1-TMR relative to P1-SiR and P1-TMR, respectively (**Figure S4**). We contextualized K_D_ with the measured fluorogenicity and included the previously published value for ATTO 655-P1-ATTO 655 ^9^ (**Figure 1C**). From this, ATTO 655 and Cy3B exhibit a decreased K_D_ (increase in binding affinity) when dual-labeled to P1 imager strand whereas SiR and TMR show the contrary. We next evaluated a shorter docking strand length to observe how K_D_ could be influenced and if it is possible for a given fluorophore to choose a strand length and sequence for an optimum binding-unbinding rate. For 8 nt P1 docking strand length, we observe an increase in K_D_ by a factor of 10 for ATTO 655-P1-ATTO 655 (**Figure S5**). Since K_D_ could also be influenced by the strand sequence itself, we repeated these experiments with the P5 imager strand (**Table S1**). P5 dual-labeled with ATTO 655 (ATTO 655-P5-ATTO 655) showed a lower K_D_ (82 nM) than ATTO 655-P1-ATTO 655 (179 nM) ^9^, while similar to single labeled P5 (P5-ATTO 655; 78 nM) (**Figure S6A**) The fluorogenicity is 4-fold for the ATTO 655-P5-ATTO 655 as compared to the 5-fold of ATTO 655-P1-ATTO 655 (**Figure S6B**).

We applied these dual-labeled, self-quenched imager strands to STED microscopy. We rationalized that the higher background signal arising from high concentrations of freely diffusing imager strands (300 nM) ^4^ would be mitigated with self-quenched imager strands. In previous work, we have shown that ATTO 655-P1-ATTO 655 reduces the fluorescence background of unbound probes, and additionally increases the fluorescence yield of bound probes ^9^. Here, we generalized this concept by investigating other fluorophores (SiR, TMR, and Cy3B) on dual-labeled imager strands with STED microscopy, enabling multi-color applications.

First, we recorded STED images of microtubules in U-2 OS cells (indirect immunolabeling for α-tubulin, see Methods) using either single-or dual-labeled P1 with SiR, TMR, and Cy3B (**Figure 2A)**. All dual-labeled imager strands exhibited a decrease in background fluorescence for unbound probes (**Figure S7A**). Interestingly, the fluorescence intensity in the STED images is different for the three fluorophores. In the case of TMR-P1-TMR and Cy3B-P1-Cy3B, the fluorescence intensity is increased, whereas SiR-P1-SiR exhibits a decrease in fluorescence intensity when bound to its complement (**Figure S7B**). Still, the background fluorescence decrease of SiR-P1-SiR relative to P1-SiR is 3-fold and the overall signal-to-background ratio (SBR) of SiR-P1-SiR is 2.3 fold higher than P1-SiR (**Figure 2B, S7C**). In conjunction, TMR-P1-TMR exhibits a SBR increase of 2.3 fold over P1-TMR and Cy3B-P1-Cy3B exhibits a SBR increase of 3.2 fold over P1-Cy3B. On initial observation, the SBR increase does not directly reflect the fluorogenicity from emission spectra but fits well when taken together with changes in K_D_, which determines the exchange of the fluorophore-labeled imager strands at the target site. SiR shows a fluorogenicity of 3.3 fold and an increase in K_D_ of 1.3 fold thereby resulting in an overall decreased SBR of 2.3 fold. TMR shows a similar behavior with a fluorogenicity of 5.2 and an increase in K_D_ of 1.4 fold and hence a decrease in SBR to 2.3 fold. Cy3B on the other hand exhibits only 1.7 fold fluorogenicity and shows a decrease in K_D_ of 0.3 fold which is observed here as an increased SBR of 3.2 fold. The overall increase in SBR also corresponds to an overall increase in spatial resolution for all dual-labeled imager strands (**Figure 2C**) (**Table S3**). Exploiting this new toolbox of dual-labeled imager strands, we performed simultaneous two-color STED imaging of mitochondria (indirect immunolabeling for TOM20) and endoplasmic reticulum (ER) (indirect immunolabeling for KDEL, see Methods) using ATTO 655-P5-ATTO 655 and TMR-P1-TMR respectively (**Figure 2D**).

**Figure 2.**
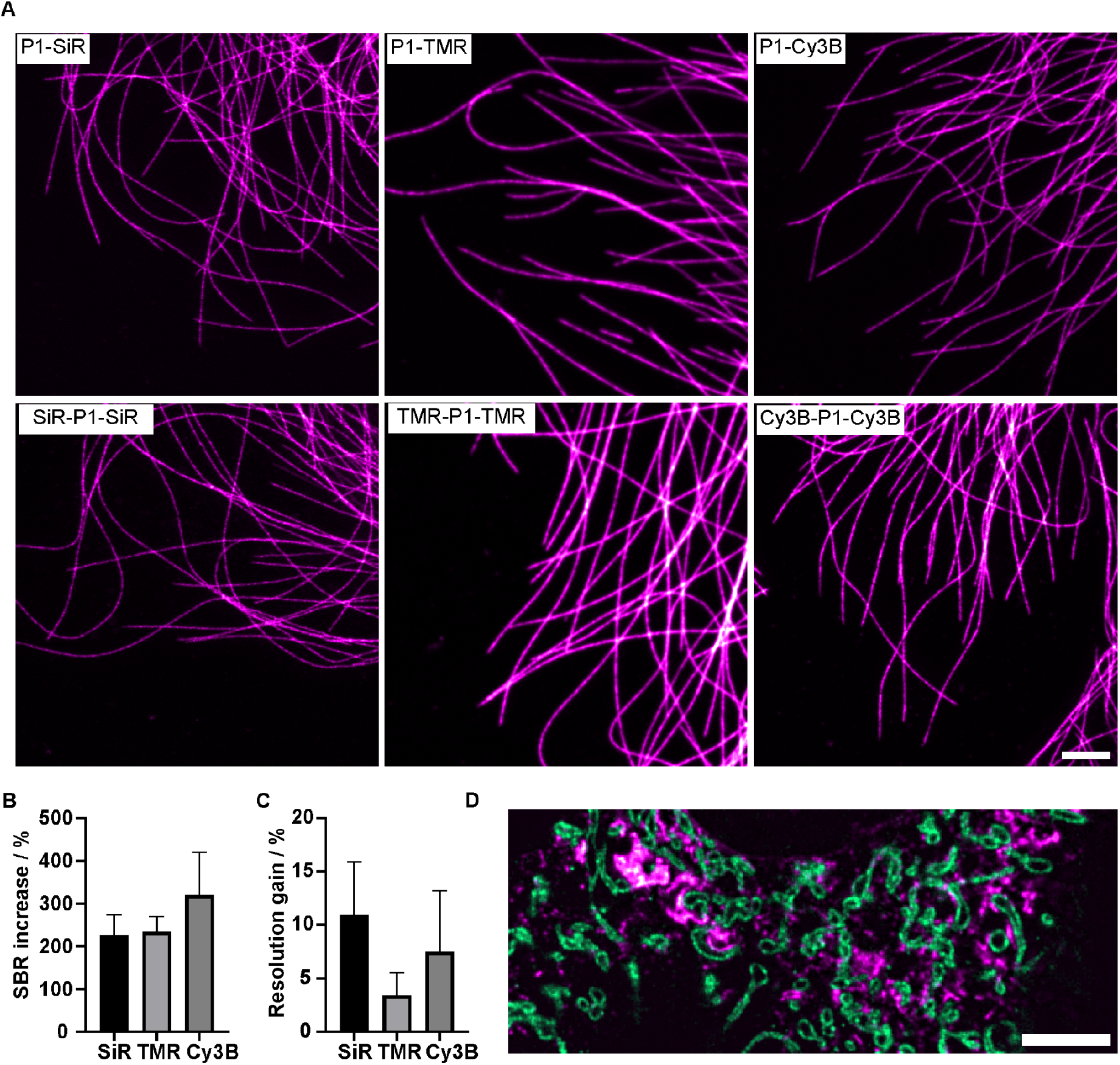
STED microscopy with single- and dual-labeled DNA oligonucleotides. **A**. STED images of U-2 OS cells immunostained for α-tubulin with different single- and dual-labeled P1 imager strands (300 nM). Scale bar is 2 μm. **B**. Percentage increase in SBR for all dual-labeled P1 imager strands over their single-labeled counterparts. **C**. Percentage increase in resolution for all dual-labeled P1 imager strands relative to single-labeled P1 imager strands. For both, B and C, n = 7 for P1-SiR and 6 for SiR-P1-SiR, For all images using single- and dual-labeled TMR and Cy3B n = 5. **D**. Dual-color STED image of a MODE-K cell immunostained for TOM20 and KDEL with ATTO 655-P5-ATTO 655 (green) and TMR-P1-TMR (magenta) respectively, scale is 5 μm.

We then explored the use of self-quenching imager strands for DNA-PAINT (**Figure 3**). We chose Cy3B and SiR for their photostability and spectral orthogonality. For Cy3B, we used 8 nt docking strands to compensate for the low K_D_ measured for 9 nt Cy3B-P1-Cy3B bound to its 9 nt complementary strand (**Figure S4**) that would be less optimal for DNA-PAINT microscopy. We used DNA-origami nanostructures as a platform for comparing single- and dual-labeled P1 imager strands. The DNA origami were constructed with three binding sites (9 nt and 8 nt docking strands for SiR and Cy3B respectively) spaced ∼ 50 nm from each other (**Figure 3A, 3C**). For both SiR and Cy3B, we determined the photon yield per target site, and we observe one population for single-labeled and two populations for dual-labeled imager strands, respectively (**Figure 3B, 3D**). For dual-labeled imager strands, the appearance of two populations can be ascribed to photobleaching of one of the fluorophores while bound to the target, or other mechanisms that deactivate one of the fluorophores (**Figure S8**). We next determined the localization precision to decrease for SiR-P1-SiR (12.3 nm) compared to P1-SiR (6.6 nm), as well as for Cy3B-P1-Cy3B (5.5 nm) and P1-Cy3B (8.0 nm) (**Figure 3B, 3D, inlays**). We attribute the larger change in localization precision for SiR to a higher background signal of P1-SiR as compared to SiR-P1-SiR (see **Figure 1Bii**). In addition, we analyzed the binding times (t_on_) of the imager strands through hybridization to the docking strands. We observed a decrease in t_on_ for SiR-P1-SiR (239 ms) over P1-SiR (410 ms) and a small change in t_on_ for Cy3B-P1-Cy3B (351 ms) over P1-Cy3B (344 ms) (**Figure S9**).

**Figure 3.**
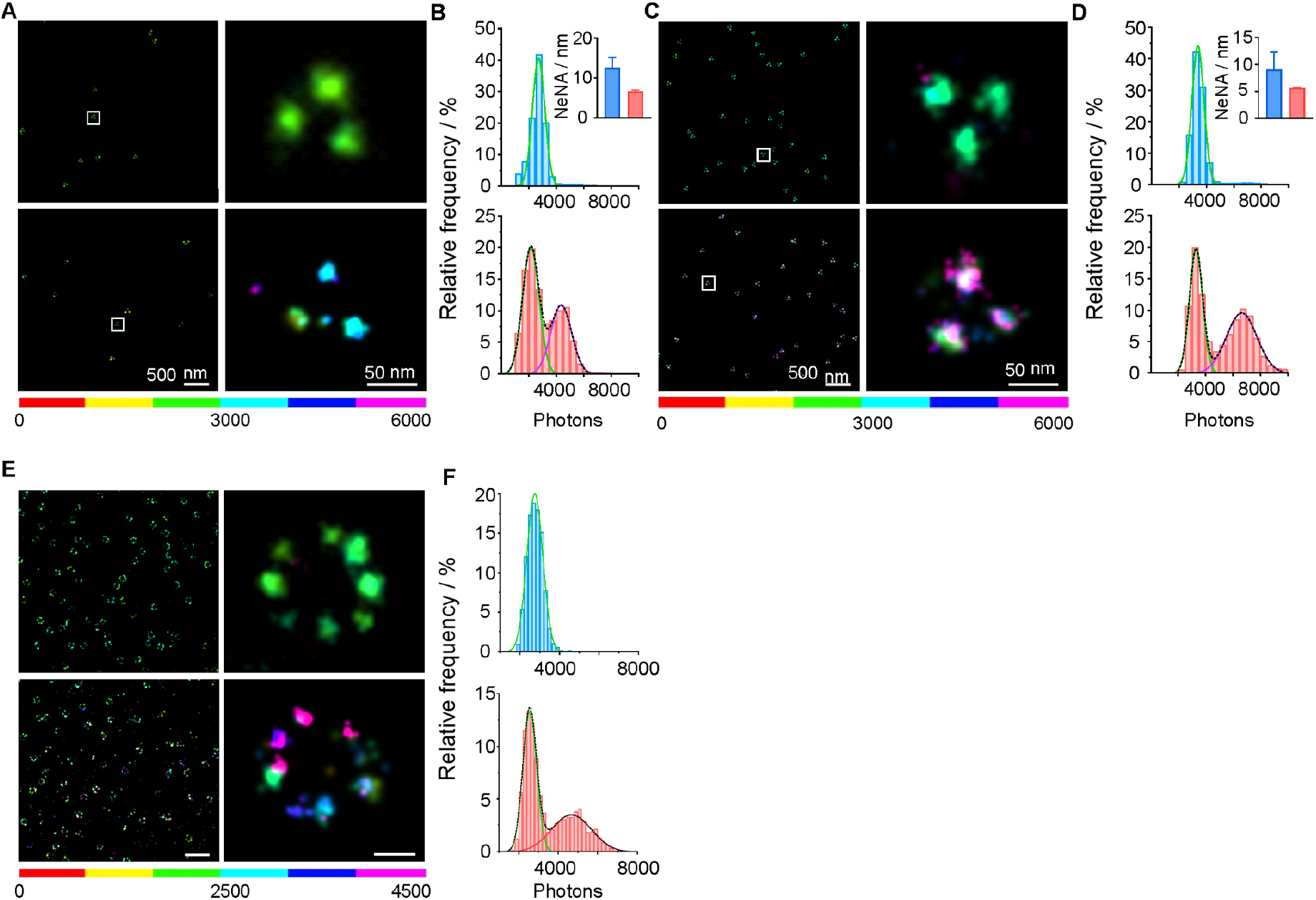
Self-quenching fluorophores for DNA-PAINT microscopy. **A**. DNA-PAINT overview image and zoom-in of DNA-origami with three binding sites (9 nt docking strands) spaced ∼ 50 nm from each other acquired with 10 nM of P1-SiR (top) and SiR-P1-SiR (bottom). **B**. Frequency distribution of photons from A acquired with P1-SiR (top, blue) and SiR-P1-SiR (bottom, red). Inlay shows the localization precision calculated using the nearest neighbor analysis (NeNA) of both overview images in A. **C**. DNA-PAINT overview image and zoom-in of DNA-origami with three binding sites (8 nt docking strands) spaced ∼ 50 nm from each other acquired with 10 nM of P1-Cy3B (top) and Cy3B-P1-Cy3B (bottom). **D**. Frequency distribution of photons from C acquired with P1-Cy3B (top, blue) and Cy3B-P1-Cy3B (bottom, red). Inlay shows the localization precision calculated using the nearest neighbor analysis (NeNA) of both overview images in C. **E**. DNA-PAINT image of nuclear pore complex protein Nup96 (NPC) and a zoomed-in image acquired using 1.5 nM of P1-Cy3B and Cy3B-P1-Cy3B. Complete DNA-PAINT overview image of NPC is shown in **Figure S10A, C. F**. Frequency distribution of photons from E acquired with P1-Cy3B (top, blue) and Cy3B-P1-Cy3B (bottom, red).

Next, we applied single- and dual-labeled imager strands for DNA-PAINT imaging in U-2 OS and visualized the nuclear pore complex (NPC) protein Nup96 (**Figure 3E, F; Figure S10**). We quantified the photon output for both P1-Cy3B and Cy3B-P1-Cy3B by choosing NPCs located in the same area of the camera field of view, in order to ensure equal illumination conditions, and exhibiting a circular structure with 8-fold symmetry. We found a two-population photon distribution (**Figure 3F**) and better localization precision in images generated with dual-labeled imager strands (7.6 nm for Cy3B-P1-Cy3B vs 10.6 nm for P1-Cy3B (**Table S4**)). This demonstrates that the extended repertoire of self-quenchable fluorophores achieve a higher photon yield and improved localization precision in DNA-PAINT microscopy.

## Discussion

We report self-quenching of fluorophore dimers attached to short DNA oligonucleotides as a general concept for DNA-based microscopy. Expanding the first reports on oxazine dimers ^9,10^, we report a fluorogenicity of DNA imager strands dual-labeled with a silicon-rhodamine (SiR), a rhodamine (TMR) or a carbocyanine (Cy3B), in a range of 1.7 to 5.2. Fluorescence quenching occurs through the formation of a short-distance H-dimer ^11^, and the fluorescence signal is restored upon hybridization to a sequence-complementary strand. This distinguishes the present concept from other reports that used e.g. Förster resonance energy transfer (FRET) which occurs at longer distances and demands for extension of DNA probes ^8,12^. A second unique property of our concept is that target binding, i.e. hybridization to a sequence-complementary DNA probe, comes with an almost two-fold increase in fluorescence intensity that is contributed by the two fluorophores. Furthermore, we observed that the hybridization affinity of dual-labeled probes is different to those of single-labeled probes, and depends on the fluorophore and the DNA sequence, which opens the possibility of further optimization.

We demonstrate the application of self-quenched DNA probes in two major super-resolution microscopy techniques, STED microscopy and DNA-PAINT microscopy. For STED microscopy, we found an increase in signal-to-background (SBR) for all dual-labeled probes as compared to single-labeled probes, ranging between 2- and 3-fold. Main parameters that determined the SBR were the fluorogenicity of the probes and the dissociation constant (K_D_). The increase in SBR was accompanied by an improved spatial resolution (**Table S3**). Further optimization is possible by adapting the hybridization kinetics to the “imaging kinetics” in STED microscopy. The availability of dual-labeled probes with orthogonal spectral properties enabled two-color STED imaging (**Figure 2E**). For DNA-PAINT microscopy, we observe an increase in photon yield, accompanied by a reduced localization precision and thus improved spatial resolution.

Beyond the application shown in this work, we envision the use of dual-labeled DNA probes for other microscopy techniques, such as SOFI ^3^ and MINFLUX ^13^. If necessary, the hybridization kinetics can be modified by sequence length and base content. These probes complement the concept of fluorogenic dimers that was reported for e.g. membrane-binding labels ^14^. An extension to more weak-affinity fluorophore labels ^15^ or protein tags ^16^ and combinations thereof ^5^ can be envisioned.

In conclusion, we show that the concept of self-quenching dimers could be extended to other classes of fluorophores, and that these probes provide a higher SBR and improved spatial resolution in cutting-edge microscopy applications.

## Acknowledgements

We thank Petra Freund for assistance with cell culture. We thank Prof. Florian Greten, Georg-Speyer-Haus, Frankfurt, for kindly providing MODE-K cells. We thank Prof. Josef Wachtveitl for access to the absorption spectrometer and Florian Hurter for experimental support. We gratefully acknowledge funding by the Deutsche Forschungsgemeinschaft (grants CRC 1177, CRC 1507 and INST 161/1020-1 FUGG) and the SubCellular Architecture of LifE (SCALE) consortium funded by the Goethe University Frankfurt, Germany.

## Methods

### DNA oligonucleotides

The sequences of the imager and docking strands used in this study are listed in **Table S1**. The sequences for DNA origami structures were purchased from Eurofins Genomics (**Table S5-S7**).

### Absorption and fluorescence spectroscopy

For absorption measurements, imager strands were diluted in 1x phosphate buffered saline (PBS, diluted from 10x stock, #14200067, Gibco, Thermo Fisher Scientific, USA), 0.5 M NaCl, 1 mM ethylenediaminetetraacetic acid (EDTA) to a final concentration of 10 μM and were analyzed in the absence and presence of docking strands (100 μM). For fluorescence spectroscopy, an imager strand concentration of 100 nM and a docking strand concentration of 100 μM were used.

For absorption spectroscopy, a Specord S600 absorption spectrometer (Analytik Jena, Germany) was used. The average was taken over 10 spectra and an integration time of 227 ms was used. Fluorescence spectra were acquired with a Cary Eclipse fluorescence spectrophotometer (Agilent Technologies, USA) with a slit size of 5 nm, mode set to slow, a gain dependent on the sample (SiR: 720;TMR: 780;Cy3B: 600) and an excitation wavelength dependent on the sample (SiR: 647 nm; TMR and Cy3B: 560 nm).

Dissociation constants for the different imager strands were determined by performing a titration series with a fixed concentration of imager strand (1 nM imager strands) and different concentrations of the complementary strand (0 - 80 μM for SiR; 0 - 40 μM for TMR; 0 - 20 μM for Cy3B) and reading out the fluorescence. A home-built confocal setup described elsewhere ^17^ was used for measuring the fluorescence. Briefly, a 647 nm (for SiR) / 532 nm (for TMR and Cy3B) continuous wave (CW) solid state laser (both from Coherent, USA) passed an acousto-optic tunable filter and was coupled into a single-mode optical fiber. The excitation light was expanded by a collimator and reflected by a dichroic mirror into a water-immersion objective (60x, 1.2 NA,UPlanSApo, Olympus, Japan). In the detection path, out-of-focus light was filtered out by a 100 μm pinhole and the fluorescence was filtered using a bandpass filter (ET 700/75 for SiR (AHF Analysentechnik, Germany) and HC 590/20 for TMR and Cy3B (AHF Analysentechnik, Germany)) which was then split using a beam splitter onto two avalanche photodiodes (APDs). The samples were warmed to 21°C by a custom-built water-cooling system and the fluorescence intensity was measured at 50 μm depth from the glass surface. A laser intensity of about 500 μW (measured at the back-focal plane of the objective) was used for excitation. For binding curves, each sample was recorded 3x for 60 s. The binding curves were measured 3x in independent experiments.

Fluorescence intensities *I* of the imager strands in dependency of the docking strand concentration *[DS]* were analyzed with OriginPro (Version 2024, OriginLab Corp., USA) by plotting the average intensity against the docking strand concentration. Binding curves were fitted with the Hill function^18^ to extract the dissociation constant *K*_*D*_:

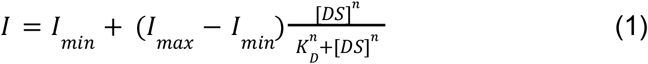

*I*_*min*_ and *I*_*max*_ are the minimum and maximum fluorescence intensities observed for the free imager strand and for the complex of imager and docking strand, respectively. The Hill coefficient *n* was set to 1 as imager and docking strand associate in a 1:1 stoichiometry.

### Fluorescence correlation spectroscopy

For FCS measurements, the above described home-built confocal setup was used.^17^ The fluorescence signal detected from two APDs was cross-correlated with a real-time correlator card (Flex03lq, correlator.com; Bridgewater, USA). 1 nM (9 nt) of fluorophore-labeled imager strand was measured in the absence and in the presence of saturating concentration of docking strand at 21°C. At this docking strand concentration, roughly all imager strands are bound to their complementary strands. The laser focus was positioned at a depth of 50 μm in the sample from the glass surface. The excitation intensity was about 500 μW at the back-focal plane of the objective. Three measurements per sample were performed, each 60 s. The correlation curves were analyzed in OriginPro. The three measurements were averaged and a three-dimensional diffusion model with a single diffusing component and either three or two or one sub-diffusional kinetics was fitted to the data.

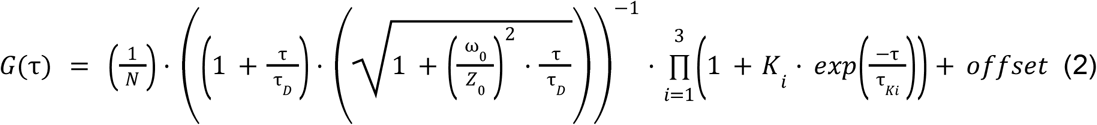

Here, *N* is the average number of emitting fluorophores in the observation volume, while τ _*D*_ is the average diffusion time through the observation volume and 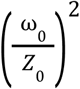 is the shape factor.

The second term describes the photophysical kinetics with *K* describing the equilibrium constant and τ_*K*_ the relaxation time of the process. For the sub-diffusional kinetics arising from 10^-7^ s the amplitude *K* can also be described in terms of the fraction D of the non-fluorescing species:

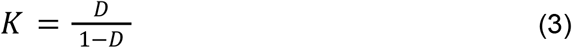

An equilibrium of the unbound imager strand between a fluorescing state F and a non-fluorescing, self-quenched species D with the rates *k*_*dark*_ and *k*_*fluorescence*_ is assumed.

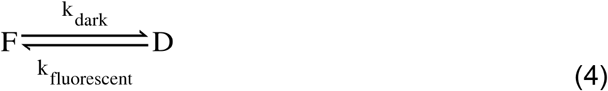

The rates of the switching between fluorescing and dark state can be described in terms of the equilibrium constant 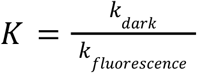 and the relaxation time 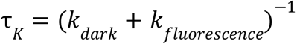^11,11^.

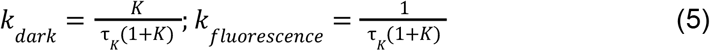

### Labeling of antibodies with DNA oligonucleotides

Unlabelled secondary antibodies (AffiniPure goat anti-mouse IgG, #115-005-003; AffiniPure donkey anti-mouse IgG, #715-005-150; AffiniPure donkey anti-rabbit IgG, #711-005-152 (Jackson ImmunoResearch, USA)) were self-labeled with azide-modified docking strands (P1 in the case of goat anti-mouse, donkey anti-mouse antibody and P5 in the case of donkey anti-rabbit antibody) according to a previously published protocol ^20^. In brief, the antibody was concentrated with an Amicon spin filter (MWCO 100 kDa, Merck, Germany) when the antibody concentration was below 2 mg/mL. The DBCO-sulfo-NHS ester linker (Jena Bioscience, Germany) was dissolved in dimethylformamide (DMF) (Sigma-Aldrich, Germany) and then diluted in 1x phosphate buffer saline (PBS). Linker and antibody were mixed in a 10:1 molar ratio and incubated for 90 min at 4°C gently shaking. The unbound linker was removed using Zeba desalting columns (Thermo Fisher Scientific, Germany). Azide-modified P1 docking strand and linker-antibody conjugate were incubated in a 10:1 molar ratio overnight at 4°C while slightly shaking. The next day, unbound DNA was removed using an Amicon spin filter (MWCO 100 kDa). The docking strand-labeled antibody was stored at 4°C.

### Cell culture

U-2 OS cells (CLS Cell Lines Service GmbH, Germany) were utilized for α-Tubulin measurements and MODE-K ^21^ cells were used for TOM20 and KDEL measurements. U-2 OS cells were cultured in Dulbecco’s modified Eagle medium (DMEM)-F12 supplemented with 10%(v/v) fetal bovine serum (FBS), 100 U/mL penicillin, and 100 μg/mL streptomycin and 1%(v/v) GlutaMAX (all reagents from Gibco, Thermo Fisher, Germany) and MODE-K cells were cultured in Dulbecco’s modified Eagle medium (DMEM) supplemented with 10%(v/v) fetal bovine serum (FBS), 100 U/mL penicillin, and 100 μg/mL streptomycin and 1%(v/v) GlutaMAX (all reagents from Gibco, Thermo Fisher, Germany). Both cell lines were incubated at 37°C and 5% CO_2_ in an automatic CO_2_ incubator (Model C150, Binder GmbH, Germany). All cells were passaged every 3-4 days or upon reaching 80% confluency.

### Immunofluorescence of microtubules

For staining microtubules, U-2 OS cells were seeded on 8-well chambered coverglass (Sarstedt, Germany) that were coated with RGD-modified poly-L-lysine-grafted polyethylene glycol (PLL-PEG-RGD, self-synthesized as described in Harwardt et al.^22^) at a density of 15000 cells per well and were incubated overnight at 37°C and 5% CO_2_. The cells were either treated with microtubule stabilization buffer (MTSB) (80 mM PIPES pH 6.8, 1 mM MgCl_2_, 5 mM EGTA, 0.5%(v/v) Triton X-100, Sigma-Aldrich, Germany) for 30 s followed by fixation with 0.5% glutaraldehyde(v/v) (Sigma-Aldrich, Germany) in MTSB for 10 min at room temperature or directly fixed using 4%(v/v) formaldehyde (Sigma-Aldrich, Germany) and 0.1%(v/v) glutaraldehyde in 1x PBS for 20 min at 37°C. The sample was washed once with 1x PBS then treated with NaBH_4_ solution (Roth, Germany) to reduce autofluorescence from glutaraldehyde followed by washing thrice with 1x PBS. The sample was then blocked using immunofluorescence staining (IF) buffer (3%(w/v) BSA, 0.1%(v/v) Triton-X 100 in 1x PBS) for 10 min after which it was incubated with primary antibody solution (anti-alpha-tubulin, #T5168, Sigma-Aldrich, Germany) in IF buffer diluted 1:500 for 1.5 h at room temperature while shaking followed by washing thrice with 1x PBS. The sample was then treated with a self-labeled secondary antibody (P1-goat anti-mouse) solution diluted 1:100 in IF buffer for 1.5 hr at room temperature followed by washing thrice with 1x PBS. The sample was then postfixed with 4%(v/v) formaldehyde for 10 min at room temperature and washed thrice with 1x PBS.

### Immunofluorescence of NPC

For staining of NPC, U2OS-Nup96-EGFP cells were seeded on 8-well chambered coverglass (Sarstedt, Germany) coated with RGD-modified poly-L-lysine-grafted polyethylene glycol (PLL-PEG-RGD, self-synthesized as described in Harwardt et al.^22^) at a density of 30,000 cells per well and were incubated 1 day at 37°C and 5% CO_2_. The cells were fixed using 4%(v/v) formaldehyde (Sigma-Aldrich, Germany) in 1x PBS for 20 min at 37°C. The fixed cells were blocked for 30 min with a commercially available buffer (antibody incubation buffer, Massive Photonics, Germany) and then incubated for 1 h with P1 docking strand modified nanobody (Massive-tag-Q anti-GFP, Massive Photonics, Germany) diluted 1:100 in antibody incubation buffer, with gentle shaking at room temperature. Afterwards, cells were washed 3x with 1x PBS, and post-fixed with 4%(v/v) formaldehyde in 1x PBS for 10 min at room temperature. After washing again 3x with 1x PBS, gold beads (Gold nanoparticles, 100 nm diameter, Product #A11-100-NPC-DIH-1-25, NanoPartz, USA) as fiducial markers were diluted 1:5 in 1x PBS and incubated for 8 min.

### Immunofluorescence of TOM20 and KDEL

For staining of TOM20 and KDEL, MODE-K cells were seeded on 8-well chambered coverglass (Sarstedt, Germany) coated with fibronectin at a density of 10,000 cells per well and were incubated 1-2 days at 37°C and 5% CO_2_. The cells were then fixed using 4%(v/v) formaldehyde (Sigma-Aldrich, Germany) in 1x PBS for 30 min at 37 °C. The washing and fixing buffer were pre-warmed to 37°C. The fixed cells were blocked and permeabilized with blocking buffer (3% BSA + 0.1 % saponin in 1x PBS) at RT with shaking for 1 h. Then they were incubated with primary antibodies (mouse anti-KDEL: 1:200 dilution in blocking buffer; rabbit anti-TOM20: 1:200 (11802-1-1AP, Proteintech, Germany)) for 2 h at RT with shaking. Afterwards, cells were washed 3x with 1x PBS, and labeled with secondary antibodies (donkey anti-mouse P1: 1:100; donkey anti-rabbit P5: 1:100) for 2 h at RT with shaking followed by washing thrice with 1x PBS. They were then post-fixed with 4%(v/v) formaldehyde in 1x PBS for 10 min at RT. The sample was stored in 1x PBS for further imaging.

### Folding and purification of DNA origami

Annealing of a rectangular DNA origami was performed in a mixture of 40 μL 1x TE (Tris-EDTA) buffer with 12.5 mM MgCl_2_ containing 10 nM scaffold strand (M13mp18; tilibit nanosystem, Germany), 100 nM core staple strands, 1 μM biotinylated staple strands, and 1 μM DNA-PAINT handles. The mixture was incubated for 5 min at 65°C and subsequently cooled to 25°C over the course of 4 hours. The DNA origami was purified using 100 kDa MWCO ultra centrifugal filter units (Amicon Ultra, Germany). The purified origami structure was stored in FoB5 buffer (5 mM Tris pH 8, EDTA pH 8, 5 mM NaCl, 5 mM MgCl_2_) at -20°C. For strand sequences see **Tables S5**-**S7**. The sequences were adapted from the Picasso design tool ^23^.

### Immobilisation of DNA origami

The channel of a glass slide (μ-Slide VI 0.5 Glass Bottom, Ibidi, Germany) was flooded with 1x PBS. Subsequently, biotinylated BSA (1 mg/mL, dissolved in 1x PBS) was added and incubated for 15 min. After 3x washing with 1x PBS, streptavidin (0.2 mg/mL, dissolved in 1x PBS) was added and incubated for 15 min. The channel was washed 3x with 1x PBS, and filled with purified DNA origami. After incubation for 20 min the channel was washed 3x with DNA-PAINT buffer (1x PBS, 500 mM NaCl). 2 pM gold beads (100 nm diameter were chosen with regards to the wavelength of the excitation laser; λ_exc_ = 647 nm, Nanopartz, USA) in DNA-PAINT buffer were added and incubated for 10 min. The channel was washed once more with DNA-PAINT buffer and the imaging buffer was filled into the channel.

### STED imaging

STED imaging was performed on an Abberior expert line microscope (Abberior Instruments, Germany) with an Olympus IX83 body (Olympus Deutschland GmbH, Germany) using a UPLXAPO 60x NA 1.42 oil immersion objective (Olympus Deutschland GmbH, Germany). For all acquired images, the imager strands were diluted to a concentration of 300 nM in DNA-PAINT buffer (1x PBS, 500 mM NaCl). For image acquisition of microtubule samples, they were excited with either a 561/640 nm pulsed excitation laser (**Table 8**) and depleted using a 775 nm pulsed laser (**Table 8**) having a 2D doughnut point spread function and with a delay of 750 ps - 8 ns. Fluorescence was collected in the spectral range of 571-630 nm in the case of 561 nm excitation and 650-760 nm in the case of 640 nm excitation using an APD. All images were acquired with a pinhole of 0.81 airy unit (AU), line accumulation/integration of 30, pixel dwell time of 5 μs and pixel size of 15 nm.

For two color image acquisition of TOM20 and KDEL, the sample was excited with a 640 nm excitation laser (10 μW at the back focal plane) and depleted using a 775 nm pulsed laser (55 mW at the back focal plane) having a 3D top-hat point spread function and with a delay of 750 ps - 8 ns. Fluorescence was collected in the spectral range of 571-630 nm (561 nm excitation) 650-760 nm (640 nm excitation) using two APDs. The images were acquired with a pinhole of 0.81 AU, line accumulation/integration of 20, pixel dwell time of 5 μs and a pixel size of 70 nm.

### STED image analysis

Fluorescence intensity from microtubules was calculated by filtering for regions with microtubules using a skeletonised binary mask. The total signal intensity of the microtubules were measured and presented as signal/μm^2^. For background calculation, regions outside a cell were selected and the signal intensity was measured and presented as signal/μm^2^. Resolution of the STED images were calculated using image decorrelation analysis ^24^ which is available as a plugin in Fiji ^25^ with a min radius of 0 and a max radius of 1, Nr of 50 and Ng of 10. All analysis was performed in Fiji.

### DNA-PAINT imaging

DNA-PAINT imaging of the cell and origami samples were carried out on a home-built widefield setup based on a Nikon Eclipse Ti microscope described before ^9^. Briefly, the excitation light was generated by a DPSS laser at 640 nm (LPX-640L-500-CSB-PPA, Oxxius S.A, France) and a DPSS laser at 561 nm (SAPPHIRE 561-300 CW CDRH, Coherent, United States) with the required excitation power controlled by an acousto-optic tunable filter (AOTFnC-400.650-TN, AA Opto Electronic, France). To clean the beam-profile the laser was coupled by a fibre collimator (60FC-4-M6.2-33, Schäfter & Kirchhoff GmbH, Germany) into a polarisation maintaining single-mode optical fibre (PMC-E-400RGB, Schäfter & Kirchhoff GmbH, Germany) and subsequently re-collimated to a FWHM diameter of 6 mm (60FC-T-4-M50L-01, Schäfter & Kirchhoff GmbH, Germany). The collinear beam was then directed through two telescope lenses (AC255-030-A-ML and AC508-150-A-ML, Thorlabs GmbH, Germany) which focused the beam onto the back focal plane of the objective (CFI Apochromat TIRF 100XC Oil, Nikon, Japan). A mirror mounted on a motorised translation stage (MTS50-Z8, Thorlabs GmbH, Germany) was used to vary the illumination angle between widefield, HILO or TIRF. The excitation light was coupled into the microscope by means of a dielectric beamsplitter (zt405/488/561/640rpc, AHF Analysentechnik, Germany) which also transmitted the emission light into the detection beam path. The axial focus was maintained using an autofocus system (Ti-PFS, Nikon) and the lateral position was adjusted using a motorised stage (Ti-S-ER, Nikon) combined with a piezo stage (Nano-Drive, MadCityLabs, USA). After spectral filtering with a red bandpass filter (700/75 ET, Chroma Technology Corp., USA) respectively with an orange bandpass filter (610/60 ET, Chroma Technology Corp., USA) the emission light was projected onto an Andor Ixon Ultra EMCCD camera (DU-897U-CS0, Andor, North Ireland). Origami and NPC samples were measured in TIRF illumination. Parameters used in DNA-PAINT imaging have been compiled together in **Table S9**.

### DNA-PAINT data analysis

DNA-PAINT data was analyzed with the Picasso software (v0.6.0) ^26^. Single-molecule localisation was performed with Picasso Localise using the following parameters: baseline 200.4 photons, sensitivity 4.32, quantum efficiency 0.95, and a min net gradient of 200,000 for NPCs, 100,000 for SiR and 250,000 for Cy3B measurements of origamis. Super-resolved images were reconstructed using Picasso Render. The image stacks for microtubules were drift corrected using redundant cross correlation (RCC) on Picasso Render with 1000-4000 frames. Origami and NPC image stacks were drift corrected using goldbeads as fiducial markers (Gold nanoparticles, 100 nm diameter, Product #A11-100-NPC-DIH-1-25, NanoPartz, USA). Localisations from DNA origami and NPCs were filtered for the ellipticity (0-0.1) and the width of the point spread function (0.8-1.2).

The photon numbers of DNA origami were extracted from single binding sites (9 nt P1-SiR: n = 1050 binding sites, 9 nt SiR-P1-SiR: n = 471 binding sites, 9 nt P1-Cy3B: n =3732 binding sites, 9 nt Cy3B-P1-Cy3B: n = 2757 binding sites). The resulting histograms were fitted with one or two gaussian functions depending on the shape of the frequency distribution. The nearest neighbor-based analysis (NeNA) localisation precision ^27^ was extracted directly from Picasso Render. In order to extract the time t_on_ of single binding events, the localisations of single docking strands were linked with a radius of 4x NeNA and 6 transient dark frames. A binding time was extracted from each binding site and a relative frequency histogram was plotted and fitted with a single log-normal fit.^6^

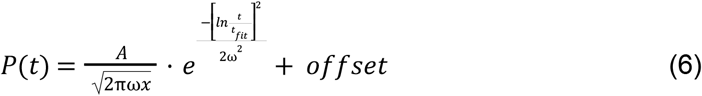

Here, *A* is the area, ω is the log standard deviation and *t*_*fit*_ is the center of the fit and the mode of the fit gives the average binding time, *t*_*on*_ of the imager strand and is given by,

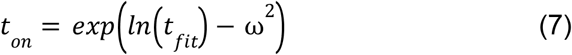

Both gaussian and log normal fits were performed in OriginPro.

