## Supplemental tables and figures for "Self-quenched fluorophore-DNA labels for super-resolution fluorescence microscopy"

[1] *Institute of Physical and Theoretical Chemistry, Goethe-University Frankfurt, Max-von-Laue-Str. 7, 60438 Frankfurt, Germany*
[2] *Optical Imaging Competence Centre, Friedrich-Alexander-Universität Erlangen-Nürnberg, Erlangen, Germany*
[#] *contributed equally*

**Supplemental Information**

### Supplemental Tables

**Table S1.** Imager and docking strands used in this study. For single-labeled imager strands, fluorophores were attached at the 3’ end. A second dye was attached to the 5’ end for the dual labeled imager strands.

| Name | Sequence | Supplier |
|---|---|---|
| 9 nt P1-SiR imager | 5’-TAGATGTAT-dye-3’ | biomers.net |
| 9 nt SiR-P1-SiR imager | 5’-dye-TAGATGTAT-dye-3’ | biomers.net |
| 9 nt P1-TMR imager | 5’-TAGATGTAT-dye-3’ | Eurofins Genomics |
| 9 nt TMR-P1-TMR imager | 5’-dye-TAGATGTAT-dye-3’ | Eurofins Genomics |
| 9 nt P1-Cy3B imager | 5’-TAGATGTAT-dye-3’ | Eurofins Genomics |
| 9 nt Cy3B-P1-Cy3B imager | 5’-dye-TAGATGTAT-dye-3’ | Eurofins Genomics |
| 9 nt P5-ATTO 655 imager | 5’-ATACATTGA-dye-3’ | Eurofins Genomics |
| 9 nt ATTO 655-P5-ATTO 655 imager | 5’-dye-ATACATTGA-dye-3’ | Eurofins Genomics |
| 9 nt ATTO 655-P1-ATTO 655 imager | 5’-dye-TAGATGTAT-dye-3’ | Eurofins Genomics |
| 9 nt P1 docking strand (spectroscopy) | 5’-TTATACATCTA-3’ | Eurofins Genomics |
| 9 nt P5 docking strand (spectroscopy) | 5’-TTTCAATGTAT-3’ | Eurofins Genomics |
| 8 nt P1 docking strand (spectroscopy) | 5’-TTATACATCT-3’ | Eurofins Genomics |
| 9 nt P1 docking strand (antibody) | 5’-azide-TTATACATCTA-3’ | Metabion |
| 9 nt P5 docking strand (antibody) | 5’-azide-TTTCAATGTAT-3’ | Metabion |
| 9 nt P1 docking strand (DNA origami) | 5’-TTTTATACATCTA-3’ | Eurofins Genomics |
| 8 nt P1 docking strand (DNA origami) | 5’-TTTTATACATCT-3’ | Eurofins Genomics |
| 9 nt P1 docking strand (nanobody) | 5’-NB-TTATACATCTA-3’ | Massive Photonics |

**Table S2**: FCS fit parameters for single- and dual-labeled imager strands unbound and saturated with docking strands.

| Sample | Diffusion time $\tau_D$ / µs | Kinetics 1 fraction | Kinetics 1 relaxation time / µs | Kinetics 2 fraction | Kinetics 2 relaxation time / µs | Quenched kinetics fraction *K* | Relaxation time $\tau_K$ of quenching / µs | Off switching rate $k_{dark}$ x$10^5$ / $s^{-1}$ | On switching rate $k_{fluorescence}$ x$10^6$ / $s^{-1}$ |
|---|---|---|---|---|---|---|---|---|---|
| P1-SiR | 826.13± 55.46 | | | 0.21 ± 0.01 | 5.69 ± 0.49 | | | | |
| P1-SiR + docking strand | 102.00± 40.20 | | | 0.23 ± 0.01 | 5.79 ± 0.36 | | | | |
| SiR-P1-SiR | 880.11 ± 32.36 | 0.11 ± 0.01 | 28.00 ± 7.21 | 0.60 ± 0.01 | 1.62 ± 0.06 | 0.21 ± 0.02 | 0.40 ± 0.05 | 4.45 ± 0.22 | 20.98 ± 2.52 |
| SiR-P1-SiR + docking strand | 1280.00 ± 84.28 | 0.13 ± 0.01 | 28.00 ± 7.21 | 0.12 ± 0.01 | 1.62 ± 0.06 | | | | |
| P1-TMR | 726.44 ± 46.50 | 0.18 ± 0.01 | 35.29 ± 11.30 | 0.24 ± 0.03 | 2.87 ± 0.26 | | | | |
| P1-TMR + docking strand | 1040.00 ± 11.44 | 0.12 ± 0.01 | 84.91 ± 6.84 | 0.29 ± 0.00 | 6.46 ± 0.25 | | | | |
| TMR-P1-TMR | 570.44 ± 69.15 | 0.28 ± 0.06 | 23.33 ± 14.57 | 0.43 ± 0.06 | 1.70 ± 0.89 | 0.38 ± 0.07 | 0.26 ± 0.14 | 12.91 ± 8.15 | 3.52 ± 2.25 |

| | | | | | | | | | |
|---|---|---|---|---|---|---|---|---|---|
| TMR-P1-TMR + docking strand | 1070.00 ± 60.58 | 0.15 ± 0.04 | 51.04 ± 27.17 | 0.26 ± 0.04 | 3.91 ± 1.07 | | | | |
| P1-Cy3B | 727.66 ± 52.05 | 0.14 ± 0.02 | 82.26 ± 12.43 | 0.10 ± 0.01 | 3.91 ± 0.46 | | | | |
| P1-Cy3B + docking strand | 844.13 ± 61.78 | 0.12 ± 0.02 | 77.54 ± 17 | 0.14 ± 0.01 | 5.30 ± 0.55 | | | | |
| Cy3B-P1-Cy3B | 870.59 ± 41.60 | 0.26 ± 0.01 | 103.90 ± 5.88 | 0.14 ± 0.04 | 4.06 ± 0.97 | 0.29 ± 0.06 | 0.32 ± 0.17 | 9.11 ± 6.50 | 2.92 ± 1.51 |
| Cy3B-P1-Cy3B + docking strand | 1160.00 ± 46.90 | 0.25 ± 0.01 | 89.44 ± 9.16 | 0.15 ± 0.01 | 3.60 ± 1.11 | | | | |

**Table S3:** Signal and background intensity, signal to background ratio (SBR) and resolution extracted from STED measurements images of microtubules with 9 nt P1 imager strands. Resolution was determined by decorrelation. Errors are given as standard deviation.

| Sample | Signal / a.u. | Background / a.u. | SBR | Resolution / nm |
|---|---|---|---|---|
| P1-SiR | 193 ± 14 | 13.8 ± 0.6 | 14.1 ± 0.9 | 91 ± 3 |
| SiR-P1-SiR | 150 ± 21 | 4.8 ± 0.4 | 32 ± 7 | 81 ± 4 |
| P1-TMR | 190 ± 30 | 4.8 ± 0.4 | 41 ± 4 | 105 ± 3 |
| TMR-P1-TMR | 250 ± 30 | 2.6 ± 0.3 | 95 ± 11 | 101 ± 3 |
| P1-Cy3B | 140 ± 30 | 6.0 ± 0.8 | 24 ± 5 | 107 ± 9 |
| Cy3B-P1-Cy3B | 290 ± 60 | 3.8 ± 0.3 | 78 ± 19 | 98 ± 4 |

**Table S4.** Localisation precision, photon numbers and binding times $t_{on}$ of P1 imager strands on DNA origami. The values were determined from DNA-PAINT measurements performed in an imaging buffer containing 1x PBS and 0.5 M NaCl. Errors are given as standard errors of the fit.

| Sample | NeNA / nm | Photon distribution first maxima | Photon distribution second maxima | $t_{on}$ / ms |
|---|---|---|---|---|
| 9 nt P1-SiR | 12.2 ± 4 | 2720 ± 18 | | 410.7 ± 1.9 |
| 9 nt SiR-P1-SiR | 6.3 ± 1.2 | 2160 ± 21 | 4370 ± 42 | 239.3 ± 7.2 |
| 9 nt P1-Cy3B | 9 ± 3 | 3385 ± 15 | | 344.7 ± 0.2 |
| 9 nt Cy3B-P1-Cy3B | 5.46 ± 0.08 | 3300 ± 20 | 6630 ± 70 | 351.1 ± 0.4 |

**Table S5.** Staple strand sequences of rectangular DNA origami.

| Position | Name | Sequence |
|---|---|---|
| A1 | 21[32]23[31]BLK | TTTTCACTCAAAGGGCGAAAAACCATCACC |
| B1 | 23[32]22[48]BLK | CAAATCAAGTTTTTTGGGGTCGAAACGTGGA |
| C1 | 21[56]23[63]BLK | AGCTGATTGCCCTTCAGAGTCCACTATTAAAGGGTGCCGT |
| D1 | 23[64]22[80]BLK | AAAGCACTAAATCGGAACCCTAATCCAGTT |

| | | |
|---|---|---|
| E1 | 21[96]23[95]BLK | AGCAAGCGTAGGGTTGAGTGTTGTAGGGAGCC |
| F1 | 23[96]22[112]BLK | CCCGATTTAGAGCTTGACGGGGAAAAAGAATA |
| G1 | 21[120]23[127]BLK | CCCAGCAGGCGAAAAATCCCTTATAAATCAAGCCGGCG |
| H1 | 21[160]22[144]BLK | TCAATATCGAACCTCAAATATCAATTCCGAAA |
| I1 | 23[128]23[159]BLK | AACGTGGCGAGAAAGGAAGGGAAACCAGTAA |
| J1 | 23[160]22[176]BLK | TAAAAGGGACATTCTGGCCAACAAAGCATC |
| K1 | 21[184]23[191]BLK | TCAACAGTTGAAAGGAGCAAATGAAAAATCTAGAGATAGA |
| L1 | 23[192]22[208]BLK | ACCCTTCTGACCTGAAAGCGTAAGACGCTGAG |
| M1 | 21[224]23[223]BLK | CTTTAGGGCCTGCAACAGTGCCAATACGTG |
| N1 | 23[224]22[240]BLK | GCACAGACAATATTTTTGAATGGGGTCAGTA |
| O1 | 21[248]23[255]BLK | AGATTAGAGCCGTCAAAAAACAGAGGTGAGGCCTATTAGT |
| P1 | 23[256]22[272]BLK | CTTTAATGCGCGAACTGATAGCCCCACCAG |
| A2 | 19[32]21[31]BLK | GTCGACTTCGGCCAACGCGCGGGGTTTTTC |
| B2 | 22[47]20[48]BLK | CTCCAACGCAGTGAGACGGGCAACCAGCTGCA |
| D2 | 22[79]20[80]BLK | TGGAACAACCGCCTGGCCCTGAGGCCCGCT |
| E2 | 19[96]21[95]BLK | CTGTGTGATTGCGTTGCGCTCACTAGAGTTGC |
| F2 | 22[111]20[112]BLK | GCCCGAGAGTCCACGCTGGTTTGCAGCTAACT |
| H2 | 19[160]20[144]BLK | GCAATTCACATATTCCTGATTATCAAAGTGTA |
| I2 | 22[143]21[159]BLK | TCGGCAAATCCTGTTTGATGGTGGACCCTCAA |
| J2 | 22[175]20[176]BLK | ACCTTGCTTGGTCAGTTGGCAAAGAGCGGA |
| L2 | 22[207]20[208]BLK | AGCCAGCAATTGAGGAAGGTTATCATCATTTT |
| M2 | 19[224]21[223]BLK | CTACCATAGTTTGAGTAACATTTAAAATAT |
| N2 | 22[239]20[240]BLK | TTAACACCAGCACTAACAACTAATCGTTATTA |
| P2 | 22[271]20[272]BLK | CAGAAGATTAGATAATACATTTGTCGACAA |
| A3 | 17[32]19[31]BLK | TGCATCTTTCCCAGTCACGACGGCCTGCAG |
| B3 | 20[47]18[48]BLK | TTAATGAACTAGAGGATCCCCGGGGGGTAACG |
| D3 | 20[79]18[80]BLK | TTCCAGTCGTAATCATGGTCATAAAAGGGG |
| E3 | 17[96]19[95]BLK | GCTTTCCGATTACGCCAGCTGGCGGCTGTTTC |
| F3 | 20[111]18[112]BLK | CACATTAAAATTGTTATCCGCTCATGCGGGCC |
| H3 | 17[160]18[144]BLK | AGAAAACAAAGAAGATGATGAAACAGGCTGCG |
| I3 | 20[143]19[159]BLK | AAGCCTGGTACGAGCCGGAAGCATAGATGATG |
| J3 | 20[175]18[176]BLK | ATTATCATTCAATATAATCCTGACAATTAC |
| L3 | 20[207]18[208]BLK | GCGGAACATCTGAATAATGGAAGGTACAAAAT |
| M3 | 17[224]19[223]BLK | CATAAATCTTTGAATACCAAGTGTTAGAAC |
| N3 | 20[239]18[240]BLK | ATTTTAAAATCAAAATTATTTGCACGGATTCG |
| P3 | 20[271]18[272]BLK | CTCGTATTAGAAATTGCGTAGATACAGTAC |
| A4 | 15[32]17[31]BLK | TAATCAGCGGATTGACCGTAATCGTAACCG |
| B4 | 18[47]16[48]BLK | CCAGGGTTGCCAGTTTGAGGGGACCCGTGGGA |
| C4 | 15[64]18[64]BLK | GTATAAGCCAACCCGTCGGATTCTGACGACAGTATCGGCCGCAAGGCG |
| D4 | 18[79]16[80]BLK | GATGTGCTTCAGGAAGATCGCACAATGTGA |
| E4 | 15[96]17[95]BLK | ATATTTTGGCTTTCATCAACATTATCCAGCCA |

| | | |
|---|---|---|
| F4 | 18[111]16[112]BLK | TCTTCGCTGCACCGCTTCTGGTGCGGCCTTCC |
| G4 | 15[128]18[128]BLK | TAAATCAAAATAATTCGCGTCTCGGAAACCAGGCAAAGGGAAGG |
| H4 | 15[160]16[144]BLK | ATCGCAAGTATGTAAATGCTGATGATAGGAAC |
| I4 | 18[143]17[159]BLK | CAACTGTTGCGCCATTCGCCATTCAAACATCA |
| J4 | 18[175]16[176]BLK | CTGAGCAAAAATTAATTACATTTTGGGTTA |
| K4 | 15[192]18[192]BLK | TCAAATATAACCTCCGGCTTAGGTAACAATTTCATTTGAAGGCGAATT |
| L4 | 18[207]16[208]BLK | CGCGCAGATTACCTTTTTTAATGGGAGAGACT |
| M4 | 15[224]17[223]BLK | CCTAAATCAAAATCATAGGTCTAAACAGTA |
| N4 | 18[239]16[240]BLK | CCTGATTGCAATATATGTGAGTGATCAATAGT |
| O4 | 15[256]18[256]BLK | GTGATAAAAAGACGCTGAGAAGAGATAACCTTGCTTCTGTTCGGGAGA |
| P4 | 18[271]16[272]BLK | CTTTTACAAAATCGTCGCTATTAGCGATAG |
| A5 | 13[32]15[31]BLK | AACGCAAAATCGATGAACGGTACCGGTTGA |
| B5 | 16[47]14[48]BLK | ACAAACGGAAAAGCCCCAAAAACACTGGAGCA |
| C5 | 13[64]15[63]BLK | TATATTTTGTCATTGCCTGAGAGTGGAAGATT |
| D5 | 16[79]14[80]BLK | GCGAGTAAAAATATTTAAATTGTTACAAAG |
| E5 | 13[96]15[95]BLK | TAGGTAAACTATTTTTGAGAGATCAAACGTTA |
| F5 | 16[111]14[112]BLK | TGTAGCCATTAAAATTCGCATTAAATGCCGGA |
| G5 | 13[128]15[127]BLK | GAGACAGCTAGCTGATAAATTAATTTTTGT |
| H5 | 13[160]14[144]BLK | GTAATAAGTTAGGCAGAGGCATTTATGATATT |
| I5 | 16[143]15[159]BLK | GCCATCAAGCTCATTTTTTAACCACAAATCCA |
| J5 | 16[175]14[176]BLK | TATAACTAACAAAGAACGCGAGAACGCCAA |
| K5 | 13[192]15[191]BLK | GTAAAGTAATCGCCATATTTAACAAAACTTTT |
| L5 | 16[207]14[208]BLK | ACCTTTTTATTTTAGTTAATTTCATAGGGCTT |
| M5 | 13[224]15[223]BLK | ACAACATGCCAACGCTCAACAGTCTTCTGA |
| N5 | 16[239]14[240]BLK | GAATTTATTTAATGGTTTGAAATATTCTTACC |
| O5 | 13[256]15[255]BLK | GTTTATCAATATGCGTTATACAAACCGACCGT |
| P5 | 16[271]14[272]BLK | CTTAGATTTAAGGCGTTAAATAAAGCCTGT |
| A6 | 11[32]13[31]BLK | AACAGTTTTGTACCAAAAACATTTTATTTC |
| B6 | 14[47]12[48]BLK | AACAAGAGGGATAAAAATTTTTAGCATAAAGC |
| C6 | 11[64]13[63]BLK | GATTTAGTCAATAAAGCCTCAGAGAACCCTCA |
| D6 | 14[79]12[80]BLK | GCTATCAGAAATGCAATGCCTGAATTAGCA |
| E6 | 11[96]13[95]BLK | AATGGTCAACAGGCAAGGCAAAGAGTAATGTG |
| F6 | 14[111]12[112]BLK | GAGGGTAGGATTCAAAAGGGTGAGACATCCAA |
| G6 | 11[128]13[127]BLK | TTTGGGGATAGTAGTAGCATTAAAAGGCCG |
| H6 | 11[160]12[144]BLK | CCAATAGCTCATCGTAGGAATCATGGCATCAA |
| I6 | 14[143]13[159]BLK | CAACCGTTTCAAATCACCATCAATTCGAGCCA |
| J6 | 14[175]12[176]BLK | CATGTAATAGAATATAAAGTACCAAGCCGT |
| K6 | 11[192]13[191]BLK | TATCCGGTCTCATCGAGAACAAGCGACAAAAG |
| L6 | 14[207]12[208]BLK | AATTGAGAATTCTGTCCAGACGACTAAACCAA |
| M6 | 11[224]13[223]BLK | GCGAACCTCCAAGAACGGGTATGACAATAA |
| N6 | 14[239]12[240]BLK | AGTATAAAGTTCAGCTAATGCAGATGTCTTTC |

| | | |
|---|---|---|
| O6 | 11[256]13[255]BLK | GCCTTAAACCAATCAATAATCGGCACGCGCCT |
| P6 | 14[271]12[272]BLK | TTAGTATCACAATAGATAAGTCCACGAGCA |
| A7 | 9[32]11[31]BLK | TTTACCCCAACATGTTTTAAATTTCCATAT |
| B7 | 12[47]10[48]BLK | TAAATCGGGATTCCCAATTCTGCGATATAATG |
| C7 | 9[64]11[63]BLK | CGGATTGCAGAGCTTAATTGCTGAAACGAGTA |
| D7 | 12[79]10[80]BLK | AAATTAAGTTGACCATTAGATACTTTTGCG |
| E7 | 9[96]11[95]BLK | CGAAAGACTTTGATAAGAGGTCATATTTCGCA |
| F7 | 12[111]10[112]BLK | TAAATCATATAACCTGTTTAGCTAACCTTTAA |
| G7 | 9[128]11[127]BLK | GCTTCAATCAGGATTAGAGAGTTATTTTCA |
| H7 | 9[160]10[144]BLK | AGAGAGAAAAAAATGAAAATAGCAAGCAAACT |
| I7 | 12[143]11[159]BLK | TTCTACTACGCGAGCTGAAAAGGTTACCGCGC |
| J7 | 12[175]10[176]BLK | TTTTATTTAAGCAAATCAGATATTTTTTGT |
| K7 | 9[192]11[191]BLK | TTAGACGGCCAAATAAGAAACGATAGAAGGCT |
| L7 | 12[207]10[208]BLK | GTACCGCAATTCTAAGAACGCGAGTATTATTT |
| M7 | 9[224]11[223]BLK | AAAGTCACAAAATAAACAGCCAGCGTTTTA |
| N7 | 12[239]10[240]BLK | CTTATCATTCCCGACTTGCGGGAGCCTAATTT |
| O7 | 9[256]11[255]BLK | GAGAGATAGAGCGTCTTTCCAGAGGTTTTGAA |
| P7 | 12[271]10[272]BLK | TGTAGAAATCAAGATTAGTTGCTCTTACCA |
| A8 | 7[32]9[31]BLK | TTTAGGACAAATGCTTTAAACAATCAGGTC |
| B8 | 10[47]8[48]BLK | CTGTAGCTTGACTATTATAGTCAGTTCATTGA |
| C8 | 7[56]9[63]BLK | ATGCAGATACATAACGGGAATCGTCATAAATAAAGCAAAG |
| D8 | 10[79]8[80]BLK | GATGGCTTATCAAAAAGATTAAGAGCGTCC |
| E8 | 7[96]9[95]BLK | TAAGAGCAAATGTTTAGACTGGATAGGAAGCC |
| F8 | 10[111]8[112]BLK | TTGCTCCTTTCAAATATCGCGTTTGAGGGGGT |
| G8 | 7[120]9[127]BLK | CGTTTACCAGACGACAAAGAAGTTTTGCCATAATTCGA |
| H8 | 7[160]8[144]BLK | TTATTACGAAGAACTGGCATGATTGCGAGAGG |
| I8 | 10[143]9[159]BLK | CCAACAGGAGCGAACCAGACCGGAGCCTTTAC |
| J8 | 10[175]8[176]BLK | TTAACGTCTAACATAAAAACAGGTAACGGA |
| K8 | 7[184]9[191]BLK | CGTAGAAAATACATACCGAGGAAACGCAATAAGAAGCGCA |
| L8 | 10[207]8[208]BLK | ATCCCAATGAGAATTAACTGAACAGTTACCAG |
| M8 | 7[224]9[223]BLK | AACGCAAAGATAGCCGAACAAACCCTGAAC |
| N8 | 10[239]8[240]BLK | GCCAGTTAGAGGGTAATTGAGCGCTTTAAGAA |
| O8 | 7[248]9[255]BLK | GTTTATTTTGTCACAATCTTACCGAAGCCCTTTAATATCA |
| P8 | 10[271]8[272]BLK | ACGCTAACACCCACAAGAATTGAAAATAGC |
| A9 | 5[32]7[31]BLK | CATCAAGTAAAACGAACTAACGAGTTGAGA |
| B9 | 8[47]6[48]BLK | ATCCCCCTATACCACATTCAACTAGAAAAATC |
| D9 | 8[79]6[80]BLK | AATACTGCCCAAAAGGAATTACGTGGCTCA |
| E9 | 5[96]7[95]BLK | TCATTCAGATGCGATTTTAAGAACAGGCATAG |
| F9 | 8[111]6[112]BLK | AATAGTAAACACTATCATAACCCTCATTGTGA |
| H9 | 5[160]6[144]BLK | GCAAGGCCTCACCAGTAGCACCATGGGCTTGA |
| I9 | 8[143]7[159]BLK | CTTTTGCAGATAAAAACCAAAATAAAGACTCC |

| | | |
|---|---|---|
| J9 | 8[175]6[176]BLK | ATACCCAACAGTATGTTAGCAAATTAGAGC |
| L9 | 8[207]6[208]BLK | AAGGAAACATAAAGGTGGCAACATTATCACCG |
| M9 | 5[224]7[223]BLK | TCAAGTTTCATTAAAGGTGAATATAAAAGA |
| N9 | 8[239]6[240]BLK | AAGTAAGCAGACACCACGGAATAATATTGACG |
| P9 | 8[271]6[272]BLK | AATAGCTATCAATAGAAAATTCAACATTCA |
| A10 | 3[32]5[31]BLK | AATACGTTTGAAAGAGGACAGACTGACCTT |
| B10 | 6[47]4[48]BLK | TACGTTAAAGTAATCTTGACAAGAACCGAACT |
| D10 | 6[79]4[80]BLK | TTATACCACCAAATCAACGTAACGAACGAG |
| E10 | 3[96]5[95]BLK | ACACTCATCCATGTTACTTAGCCGAAAGCTGC |
| F10 | 6[111]4[112]BLK | ATTACCTTTGAATAAGGCTTGCCCAAATCCGC |
| H10 | 3[160]4[144]BLK | TTGACAGGCCACCACCAGAGCCGCGATTTGTA |
| I10 | 6[143]5[159]BLK | GATGGTTTGAACGAGTAGTAAATTTACCATTA |
| J10 | 6[175]4[176]BLK | CAGCAAAAGGAAACGTCACCAATGAGCCGC |
| L10 | 6[207]4[208]BLK | TCACCGACGCACCGTAATCAGTAGCAGAACCG |
| M10 | 3[224]5[223]BLK | TTAAAGCCAGAGCCGCCACCCTCGACAGAA |
| N10 | 6[239]4[240]BLK | GAAATTATTGCCTTTAGCGTCAGACCGGAACC |
| P10 | 6[271]4[272]BLK | ACCGATTGTCGGCATTTTCGGTCATAATCA |
| A11 | 1[32]3[31]BLK | AGGCTCCAGAGGCTTTGAGGACACGGGTAA |
| B11 | 4[47]2[48]BLK | GACCAACTAATGCCACTACGAAGGGGGTAGCA |
| C11 | 1[64]4[64]BLK | TTTATCAGGACAGCATCGGAACGACACCAACCTAAAACGAGGTCAATC |
| D11 | 4[79]2[80]BLK | GCGCAGACAAGAGGCAAAAGAATCCCTCAG |
| E11 | 1[96]3[95]BLK | AAACAGCTTTTTGCGGGATCGTCAACACTAAA |
| F11 | 4[111]2[112]BLK | GACCTGCTCTTTGACCCCCAGCGAGGGAGTTA |
| G11 | 1[128]4[128]BLK | TGACAACTCGCTGAGGCTTGCATTATACCAAGCGCGATGATAAA |
| H11 | 1[160]2[144]BLK | TTAGGATTGGCTGAGACTCCTCAATAACCGAT |
| I11 | 4[143]3[159]BLK | TCATCGCCAACAAAGTACAACGGACGCCAGCA |
| J11 | 4[175]2[176]BLK | CACCAGAAAGGTTGAGGCAGGTCATGAAAG |
| K11 | 1[192]4[192]BLK | GCGGATAACCTATTATTCTGAAACAGACGATTGGCCTTGAAGAGCCAC |
| L11 | 4[207]2[208]BLK | CCACCCTCTATTCACAAACAAATACCTGCCTA |
| M11 | 1[224]3[223]BLK | GTATAGCAAACAGTTAATGCCCAATCCTCA |
| N11 | 4[239]2[240]BLK | GCCTCCCTCAGAATGGAAAGCGCAGTAACAGT |
| O11 | 1[256]4[256]BLK | CAGGAGGTGGGGTCAGTGCCTTGAGTCTCTGAATTTACCGGGAACCAG |
| P11 | 4[271]2[272]BLK | AAATCACCTTCCAGTAAGCGTCAGTAATAA |
| A12 | 0[47]1[31]BLK | AGAAAGGAACAACTAAAGGAATTCAAAAAAA |
| B12 | 2[47]0[48]BLK | ACGGCTACAAAAGGAGCCTTTAATGTGAGAAT |
| C12 | 0[79]1[63]BLK | ACAACTTTCAACAGTTTCAGCGGATGTATCGG |
| D12 | 2[79]0[80]BLK | CAGCGAAACTTGCTTTCGAGGTGTTGCTAA |
| E12 | 0[111]1[95]BLK | TAAATGAATTTTCTGTATGGGATTAATTTCTT |
| F12 | 2[111]0[112]BLK | AAGGCCGCTGATACCGATAGTTGCGACGTTAG |
| G12 | 0[143]1[127]BLK | TCTAAAGTTTTGTCGTCTTTCCAGCCGACAA |
| H12 | 0[175]0[144]BLK | TCCACAGACAGCCCTCATAGTTAGCGTAACGA |

| | | |
|---|---|---|
| I12 | 2[143]1[159]BLK | ATATTCGGAACCATCGCCCACGCAGAGAAGGA |
| J12 | 2[175]0[176]BLK | TATTAAGAAGCGGGGTTTTGCTCGTAGCAT |
| K12 | 0[207]1[191]BLK | TCACCAGTACAAACTACAACGCCTAGTACCAG |
| L12 | 2[207]0[208]BLK | TTTCGGAAGTGCCGTCGAGAGGGTGAGTTTCG |
| M12 | 0[239]1[223]BLK | AGGAACCCATGTACCGTAACACTTGATATAA |
| N12 | 2[239]0[240]BLK | GCCCGTATCCGGAATAGGTGTATCAGCCCAAT |
| O12 | 0[271]1[255]BLK | CCACCCTCATTTTCAGGGATAGCAACCGTACT |
| P12 | 2[271]0[272]BLK | GTTTTAACTTAGTACCGCCACCCAGAGCCA |

**Table S6.** Sequences of biotinylated staple strands for DNA origami.

| Position | Name | Sequence | Modification |
|---|---|---|---|
| C2 | 18[63]20[56]BIOTIN | ATTAAGTTTACCGAGCTCGAATTCGGGAAACCTGTCGTGC | 5'-biotin |
| C9 | 4[63]6[56]BIOTIN | ATAAGGGAACCGGATATTCATTACGTCAGGACGTTGGGAA | 5'-biotin |
| G2 | 18[127]20[120]BIOTIN | GCGATCGGCAATTCCACACAACAGGTGCCTAATGAGTG | 5'-biotin |
| G9 | 4[127]6[120]BIOTIN | TTGTGTCGTGACGAGAAACACCAAATTTCAACTTTAAT | 5'-biotin |
| K2 | 18[191]20[184]BIOTIN | ATTCATTTTTGTTTGGATTATACTAAGAAACCACCAGAAG | 5'-biotin |
| K9 | 4[191]6[184]BIOTIN | CACCCTCAGAAACCATCGATAGCATTGAGCCATTTGGGAA | 5'-biotin |
| O2 | 18[255]20[248]BIOTIN | AACAATAACGTAAAACAGAAATAAAAATCCTTTGCCCGAA | 5'-biotin |
| O9 | 4[255]6[248]BIOTIN | AGCCACCACTGTAGCGCGTTTTCAAGGGAGGGAAGGTAAA | 5'-biotin |

**Table S7.** Sequence of scaffold strand (M13mp18) for DNA origami.

AATGCTACTACTATTAGTAGAATTGATGCCACCTTTTCAGCTCGCGCCCCAAATGAAAATATAGCTAAACAGGTTATT
GACCATTTGCGAAATGTATCTAATGGTCAAACTAAATCTACTCGTTCGCAGAATTGGGAATCAACTGTTATATGGAAT
GAAACTTCCAGACACCGTACTTTAGTTGCATATTTAAAACATGTTGAGCTACAGCATTATATTCAGCAATTAAGCTCT
AAGCCATCCGCAAAAATGACCTCTTATCAAAAGGAGCAATTAAAGGTACTCTCTAATCCTGACCTGTTGGAGTTTG
CTTCCGGTCTGGTTCGCTTTGAAGCTCGAATTAAAACGCGATATTTGAAGTCTTTCGGGCTTCCTCTTAATCTTTTT
GATGCAATCCGCTTTGCTTCTGACTATAATAGTCAGGGTAAAGACCTGATTTTTGATTTATGGTCATTCTCGTTTTCT
GAACTGTTTAAAGCATTTGAGGGGGATTCAATGAATATTTATGACGATTCCGCAGTATTGGACGCTATCCAGTCTAA
ACATTTTACTATTACCCCCTCTGGCAAAACTTCTTTTGCAAAAGCCTCTCGCTATTTTGGTTTTTATCGTCGTCTGGT
AAACGAGGGTTATGATAGTGTTGCTCTTACTATGCCTCGTAATTCCTTTTGGCGTTATGTATCTGCATTAGTTGAATG
TGGTATTCCTAAATCTCAACTGATGAATCTTTCTACCTGTAATAATGTTGTTCCGTTAGTTCGTTTTATTAACGTAGAT
TTTTCTTCCCAACGTCCTGACTGGTATAATGAGCCAGTTCTTAAAATCGCATAAGGTAATTCACAATGATTAAAGTTG
AAATTAAACCATCTCAAGCCCAATTTACTACTCGTTCTGGTGTTTCTCGTCAGGGCAAGCCTTATTCACTGAATGAG
CAGCTTTGTTACGTTGATTTGGGTAATGAATATCCGGTTCTTGTCAAGATTACTCTTGATGAAGGTCAGCCAGCCTA
TGCGCCTGGTCTGTACACCGTTCATCTGTCCTCTTTCAAAGTTGGTCAGTTCGGTTCCCTTATGATTGACCGTCTG
CGCCTCGTTCCGGCTAAGTAACATGGAGCAGGTCGCGGATTTCGACACAATTTATCAGGCGATGATACAAATCTCC
GTTGTACTTTGTTTCGCGCTTGGTATAATCGCTGGGGGTCAAAGATGAGTGTTTTAGTGTATTCTTTTGCCTCTTTC
GTTTTAGGTTGGTGCCTTCGTAGTGGCATTACGTATTTTACCCGTTTAATGGAAACTTCCTCATGAAAAAGTCTTTA
GTCCTCAAAGCCTCTGTAGCCGTTGCTACCCTCGTTCCGATGCTGTCTTTCGCTGCTGAGGGTGACGATCCCGC
AAAAGCGGCCTTTAACTCCCTGCAAGCCTCAGCGACCGAATATATCGGTTATGCGTGGGCGATGGTTGTTGTCATT
GTCGGCGCAACTATCGGTATCAAGCTGTTTAAGAAATTCACCTCGAAAGCAAGCTGATAAACCGATACAATTAAAG
GCTCCTTTTGGAGCCTTTTTTTTGGAGATTTTCAACGTGAAAAAATTATTATTCGCAATTCCTTTAGTTGTTCCTTTC
TATTCTCACTCCGCTGAAACTGTTGAAAGTTGTTTAGCAAAATCCCATACAGAAAATTCATTTACTAACGTCTGGAA
AGACGACAAAACTTTAGATCGTTACGCTAACTATGAGGGCTGTCTGTGGAATGCTACAGGCGTTGTAGTTTGTACT
GGTGACGAAACTCAGTGTTACGGTACATGGGTTCCTATTGGGCTTGCTATCCCTGAAAATGAGGGTGGTGGCTCT

GAGGGTGGCGGTTCTGAGGGTGGCGGTTCTGAGGGTGGCGGTACTAAACCTCCTGAGTACGGTGATACACCTAT
TCCGGGCTATACTTATATCAACCCTCTCGACGGCACTTATCCGCCTGGTACTGAGCAAAACCCCGCTAATCCTAAT
CCTTCTCTTGAGGAGTCTCAGCCTCTTAATACTTTCATGTTTCAGAATAATAGGTTCCGAAATAGGCAGGGGGCATT
AACTGTTTATACGGGCACTGTTACTCAAGGCACTGACCCCGTTAAAACTTATTACCAGTACACTCCTGTATCATCAA
AAGCCATGTATGACGCTTACTGGAACGGTAAATTCAGAGACTGCGCTTTCCATTCTGGCTTTAATGAGGATTTATTT
GTTTGTGAATATCAAGGCCAATCGTCTGACCTGCCTCAACCTCCTGTCAATGCTGGCGGCGGCTCTGGTGGTGG
TTCTGGTGGCGGCTCTGAGGGTGGTGGCTCTGAGGGTGGCGGTTCTGAGGGTGGCGGCTCTGAGGGAGGCGG
TTCCGGTGGTGGCTCTGGTTCCGGTGATTTTGATTATGAAAAGATGGCAAACGCTAATAAGGGGGCTATGACCGA
AAATGCCGATGAAAACGCGCTACAGTCTGACGCTAAAGGCAAACTTGATTCTGTCGCTACTGATTACGGTGCTGC
TATCGATGGTTTCATTGGTGACGTTTCCGGCCTTGCTAATGGTAATGGTGCTACTGGTGATTTTGCTGGCTCTAATT
CCCAAATGGCTCAAGTCGGTGACGGTGATAATTCACCTTTAATGAATAATTTCCGTCAATATTTACCTTCCCTCCCT
CAATCGGTTGAATGTCGCCCTTTTGTCTTTGGCGCTGGTAAACCATATGAATTTTCTATTGATTGTGACAAAATAAAC
TTATTCCGTGGTGTCTTTGCGTTTCTTTTATATGTTGCCACCTTTATGTATGTATTTTCTACGTTTGCTAACATACTGC
GTAATAAGGAGTCTTAATCATGCCAGTTCTTTTGGGTATTCCGTTATTATTGCGTTTCCTCGGTTTCCTTCTGGTAAC
TTTGTTCGGCTATCTGCTTACTTTTCTTAAAAAGGGCTTCGGTAAGATAGCTATTGCTATTTCATTGTTTCTTGCTCT
TATTATTGGGCTTAACTCAATTCTTGTGGGTTATCTCTCTGATATTAGCGCTCAATTACCCTCTGACTTTGTTCAGGG
TGTTCAGTTAATTCTCCCGTCTAATGCGCTTCCCTGTTTTTATGTTATTCTCTCTGTAAAGGCTGCTATTTTCATTTTT
GACGTTAAACAAAAAATCGTTTCTTATTTGGATTGGGATAAATAATATGGCTGTTTATTTTGTAACTGGCAAATTAGG
CTCTGGAAAGACGCTCGTTAGCGTTGGTAAGATTCAGGATAAAATTGTAGCTGGGTGCAAAATAGCAACTAATCTT
GATTTAAGGCTTCAAAACCTCCCGCAAGTCGGGAGGTTCGCTAAAACGCCTCGCGTTCTTAGAATACCGGATAAG
CCTTCTATATCTGATTTGCTTGCTATTGGGCGCGGTAATGATTCCTACGATGAAAATAAAAACGGCTTGCTTGTTCT
CGATGAGTGCGGTACTTGGTTTAATACCCGTTCTTGGAATGATAAGGAAAGACAGCCGATTATTGATTGGTTTCTAC
ATGCTCGTAAATTAGGATGGGATATTATTTTTCTTGTTCAGGACTTATCTATTGTTGATAAACAGGCGCGTTCTGCAT
TAGCTGAACATGTTGTTTATTGTCGTCGTCTGGACAGAATTACTTTACCTTTTGTCGGTACTTTATATTCTCTTATTAC
TGGCTCGAAAATGCCTCTGCCTAAATTACATGTTGGCGTTGTTAAATATGGCGATTCTCAATTAAGCCCTACTGTTG
AGCGTTGGCTTTATACTGGTAAGAATTTGTATAACGCATATGATACTAAACAGGCTTTTTCTAGTAATTATGATTCCGG
TGTTTATTCTTATTTAACGCCTTATTTATCACACGGTCGGTATTTCAAACCATTAAATTTAGGTCAGAAGATGAAATTA
ACTAAAATATATTTGAAAAAGTTTTCTCGCGTTCTTTGTCTTGCGATTGGATTTGCATCAGCATTTACATATAGTTATAT
AACCCAACCTAAGCCGGAGGTTAAAAAGGTAGTCTCTCAGACCTATGATTTTGATAAATTCACTATTGACTCTTCTC
AGCGTCTTAATCTAAGCTATCGCTATGTTTTCAAGGATTCTAAGGGAAAATTAATTAATAGCGACGATTTACAGAAGC
AAGGTTATTCACTCACATATATTGATTTATGTACTGTTTCCATTAAAAAAGGTAATTCAAATGAAATTGTTAAATGTAAT
TAATTTTGTTTTCTTGATGTTTGTTTCATCATCTTCTTTTGCTCAGGTAATTGAAATGAATAATTCGCCTCTGCGCGAT
TTTGTAACTTGGTATTCAAAGCAATCAGGCGAATCCGTTATTGTTTCTCCCGATGTAAAAGGTACTGTTACTGTATAT
TCATCTGACGTTAAACCTGAAAATCTACGCAATTTCTTTATTTCTGTTTTACGTGCAAATAATTTTGATATGGTAGGTT
CTAACCCTTCCATTATTCAGAAGTATAATCCAAACAATCAGGATTATATTGATGAATTGCCATCATCTGATAATCAGGA
ATATGATGATAATTCCGCTCCTTCTGGTGGTTTCTTTGTTCCGCAAAATGATAATGTTACTCAAACTTTTAAAATTAAT
AACGTTCGGGCAAAGGATTTAATACGAGTTGTCGAATTGTTTGTAAAGTCTAATACTTCTAAATCCTCAAATGTATTA
TCTATTGACGGCTCTAATCTATTAGTTGTTAGTGCTCCTAAAGATATTTTAGATAACCTTCCTCAATTCCTTTCAACTG
TTGATTTGCCAACTGACCAGATATTGATTGAGGGTTTGATATTTGAGGTTCAGCAAGGTGATGCTTTAGATTTTTCAT
TTGCTGCTGGCTCTCAGCGTGGCACTGTTGCAGGCGGTGTTAATACTGACCGCCTCACCTCTGTTTTATCTTCTG
CTGGTGGTTCGTTCGGTATTTTTAATGGCGATGTTTTAGGGCTATCAGTTCGCGCATTAAAGACTAATAGCCATTCA
AAAATATTGTCTGTGCCACGTATTCTTACGCTTTCAGGTCAGAAGGGTTCTATCTCTGTTGGCCAGAATGTCCCTTT
TATTACTGGTCGTGTGACTGGTGAATCTGCCAATGTAAATAATCCATTTCAGACGATTGAGCGTCAAAATGTAGGTA
TTTCCATGAGCGTTTTTCCTGTTGCAATGGCTGGCGGTAATATTGTTCTGGATATTACCAGCAAGGCCGATAGTTTG
AGTTCTTCTACTCAGGCAAGTGATGTTATTACTAATCAAAGAAGTATTGCTACAACGGTTAATTTGCGTGATGGACA
GACTCTTTTACTCGGTGGCCTCACTGATTATAAAAACACTTCTCAGGATTCTGGCGTACCGTTCCTGTCTAAAATCC
CTTTAATCGGCCTCCTGTTTAGCTCCCGCTCTGATTCTAACGAGGAAAGCACGTTATACGTGCTCGTCAAAGCAAC
CATAGTACGCGCCCTGTAGCGGCGCATTAAGCGCGGCGGGTGTGGTGGTTACGCGCAGCGTGACCGCTACACTT
GCCAGCGCCCTAGCGCCCGCTCCTTTCGCTTTCTTCCCTTCCTTTCTCGCCACGTTCGCCGGCTTTCCCCGTCA
AGCTCTAAATCGGGGGCTCCCTTTAGGGTTCCGATTTAGTGCTTTACGGCACCTCGACCCCAAAAAACTTGATTTG
GGTGATGGTTCACGTAGTGGGCCATCGCCCTGATAGACGGTTTTTCGCCCTTTGACGTTGGAGTCCACGTTCTTT
AATAGTGGACTCTTGTTCCAAACTGGAACAACACTCAACCCTATCTCGGGCTATTCTTTTGATTTATAAGGGATTTT
GCCGATTTCGGAACCACCATCAAACAGGATTTTCGCCTGCTGGGGCAAACCAGCGTGGACCGCTTGCTGCAACT
CTCTCAGGGCCAGGCGGTGAAGGGCAATCAGCTGTTGCCCGTCTCACTGGTGAAAAGAAAAACCACCCTGGCG

CCCAATACGCAAACCGCCTCTCCCCGCGCGTTGGCCGATTCATTAATGCAGCTGGCACGACAGGTTTCCCGACT
GGAAAGCGGGCAGTGAGCGCAACGCAATTAATGTGAGTTAGCTCACTCATTAGGCACCCCAGGCTTTACACTTTA
TGCTTCCGGCTCGTATGTTGTGTGGAATTGTGAGCGGATAACAATTTCACACAGGAAACAGCTATGACCATGATTA
CGAATTCGAGCTCGGTACCCGGGGATCCTCTAGAGTCGACCTGCAGGCATGCAAGCTTGGCACTGGCCGTCGTT
TTACAACGTCGTGACTGGGAAAACCCTGGCGTTACCCAACTTAATCGCCTTGCAGCACATCCCCCTTTCGCCAGC
TGGCGTAATAGCGAAGAGGCCCGCACCGATCGCCCTTCCCAACAGTTGCGCAGCCTGAATGGCGAATGGCGCTT
TGCCTGGTTTCCGGCACCAGAAGCGGTGCCGGAAAGCTGGCTGGAGTGCGATCTTCCTGAGGCCGATACTGTC
GTCGTCCCCTCAAACTGGCAGATGCACGGTTACGATGCGCCCATCTACACCAACGTGACCTATCCCATTACGGTC
AATCCGCCGTTTGTTCCCACGGAGAATCCGACGGGTTGTTACTCGCTCACATTTAATGTTGATGAAAGCTGGCTAC
AGGAAGGCCAGACGCGAATTATTTTTGATGGCGTTCCTATTGGTTAAAAAATGAGCTGATTTAACAAAAATTTAATG
CGAATTTTAACAAAATATTAACGTTTACAATTTAAATATTTGCTTATACAATCTTCCTGTTTTTGGGGCTTTTCTGATTA
TCAACCGGGGTACATATGATTGACATGCTAGTTTTACGATTACCGTTCATCGATTCTCTTGTTTGCTCCAGACTCTC
AGGCAATGACCTGATAGCCTTTGTAGATCTCTCAAAAATAGCTACCCTCTCCGGCATTAATTTATCAGCTAGAACGG
TTGAATATCATATTGATGGTGATTTGACTGTCTCCGGCCTTTCTCACCCTTTTGAATCTTTACCTACACATTACTCAG
GCATTGCATTTAAAATATATGAGGGTTCTAAAAATTTTTATCCTTGCGTTGAAATAAAGGCTTCTCCCGCAAAAGTAT
TACAGGGTCATAATGTTTTTGGTACAACCGATTTAGCTTTATGCTCTGAGGCTTTATTGCTTAATTTTGCTAATTCTTT
GCCTTGCCTGTATGATTTATTGGATGTT

**Table S8**: Laser power (measured at the back focal plane) used for excitation and depletion laser for different fluorophores used in STED-PAINT imaging

| Fluorophore | Excitation laser power (µW) | Depletion laser power (mW) [775 nm] |
|---|---|---|
| SiR | 12.2 [640 nm] | 127.9 |
| TMR | 1.3 [561 nm] | 105.6 |
| Cy3B | 1.3 [561 nm] | 160.8 |

**Table S9**. Measurement parameters for DNA-PAINT imaging.

| Sample | Buffer | Imager conc. (nM) | Laser intensity (W/cm²) | Preamp gain | EM gain | Camera exposure time (ms) | Readout mode (MHz) | Frame transfer | Total measured frames |
|---|---|---|---|---|---|---|---|---|---|
| DNA origami, SiR | 1x PBS, 0.5 M NaCl | 10 | 120 (640 nm) | 3 | 200 | 80 | 5000 | on | 10000 |
| DNA origami, Cy3B | 1x PBS, 0.5 M NaCl, PCA, PCD, Trolox | 10 | 130 (571 nm) | 3 | 200 | 80 | 5000 | on | 20000 |
| Nup96 (NPC) | 1x PBS, 0.5 M NaCl, 1 mM EDTA, PCA, PCD, Trolox | 1.5 | 200 (571 nm) | 3 | 200 | 80 | 5000 | on | 40000 |

### Supplemental Figures

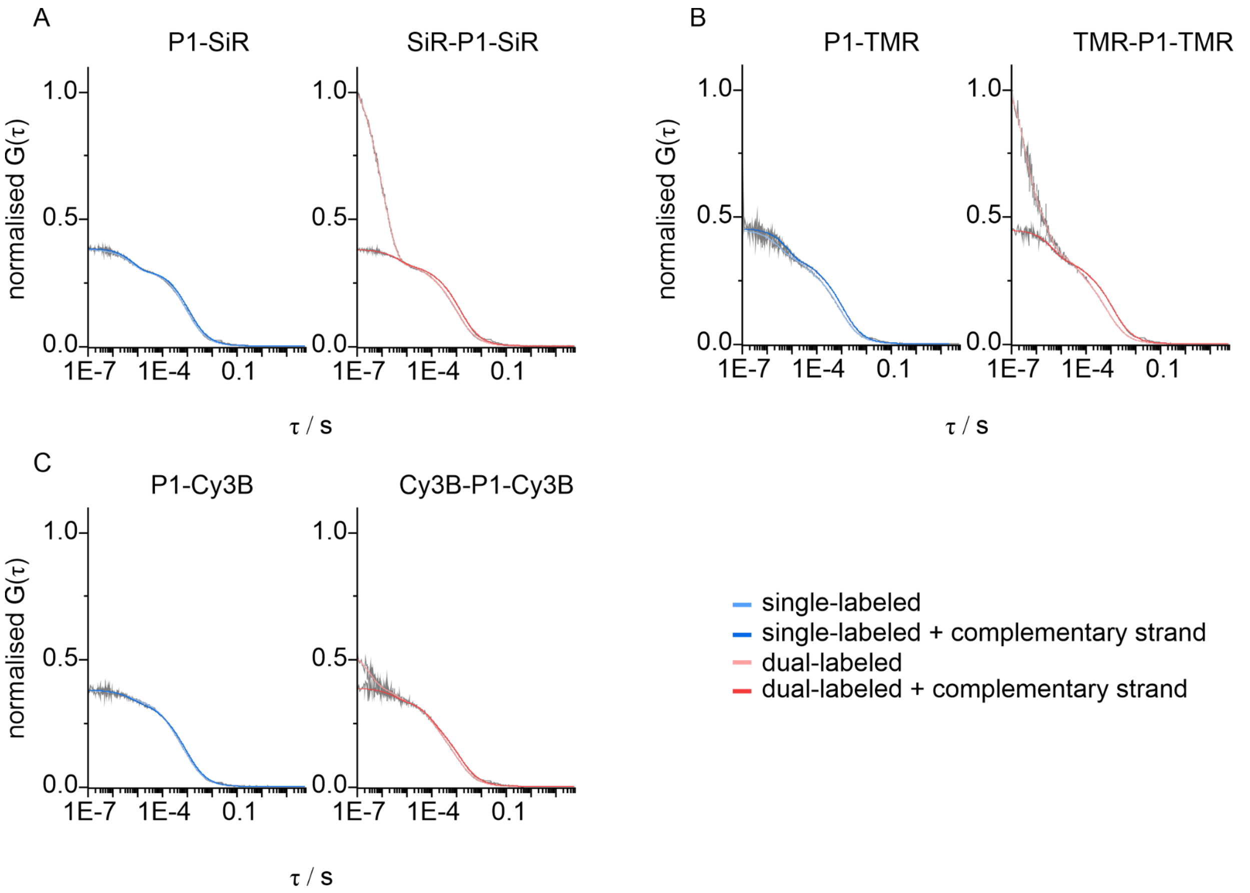


**Figure S1**: Normalised FCS cross-correlation functions (G(τ)) of single- and dual-labeled P1 imager strands with different fluorophores. **A**. G(τ) (gray) and their fits (blue/red) of P1-SiR and SiR-P1-SiR single-stranded and hybridized to its complementary docking strand. **B**. G(τ) of P1-TMR and TMR-P1-TMR single-stranded and hybridized to its complementary docking strand. **C**. G(τ) of P1-Cy3B and Cy3B-P1-Cy3B single-stranded and hybridized to its complementary docking strand. Cross correlation functions were first normalized to the number of molecules for each measurement, and then all measurements were treated as one dataset and normalized to a maximum value of 1 and a minimum value of 0.

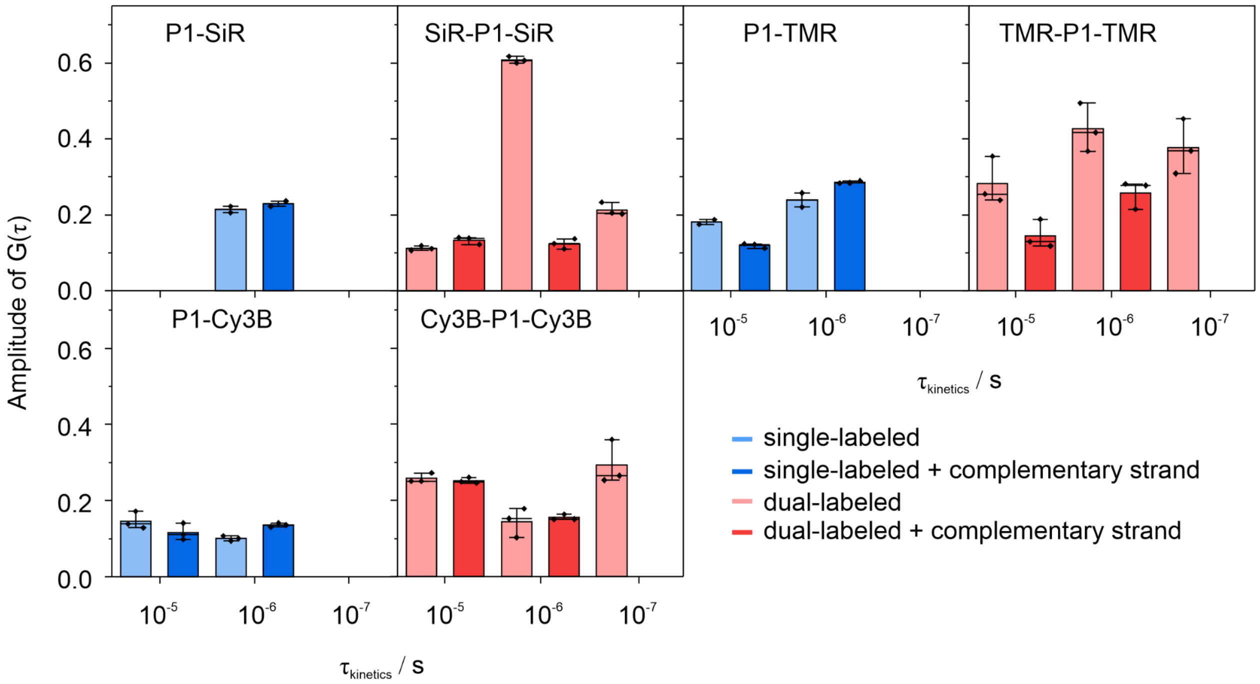


**Figure S2**: Sub-diffusional kinetics observed in single- and dual-labeled P1 imager strands with different fluorophores. Sub-diffusional kinetic amplitudes derived from G(τ) fits of single-stranded (light colored) and those hybridized to their complementary docking strands (dark colored) from single- and dual-labeled P1 imager strands with SiR, TMR and Cy3B. The amplitudes were grouped based on their time range into $10^{-5}$ , $10^{-6}$, and $10^{-7}$ s. Errors in the graphs represent standard deviation.

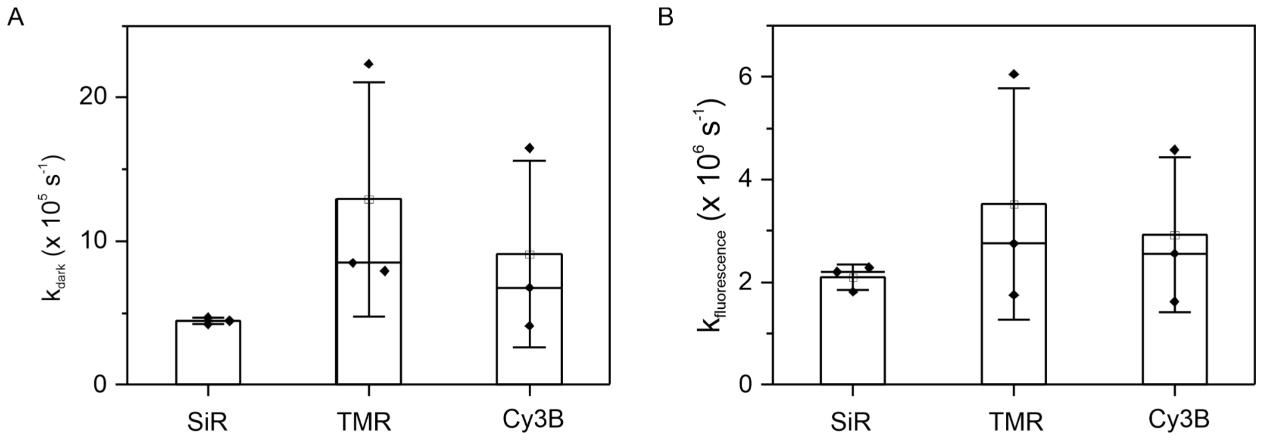


**Figure S3**: Kinetic rates of self-quenching for dual-labeled imager strands with different fluorophores. **A**. Kinetic rate of switching to a dark state in the self-quenching process ($k_{dark}$) for dual-labeled imager strands with SiR, TMR and Cy3B when single-stranded in solution. **B**. Kinetic rate of switching to a fluorescent state in the self-quenching process ($k_{fluorescence}$) for dual-labeled imager strands with SiR, TMR and Cy3B when single-stranded in solution. Both $k_{dark}$ and $k_{fluorescence}$ were calculated from the amplitude (K) and time ($\tau_k$) of the fast sub-diffusional kinetics component ($10^{-7}$s, see **Figure S2**) (**Methods equation (5)**). Errors in the graphs represent standard deviation.

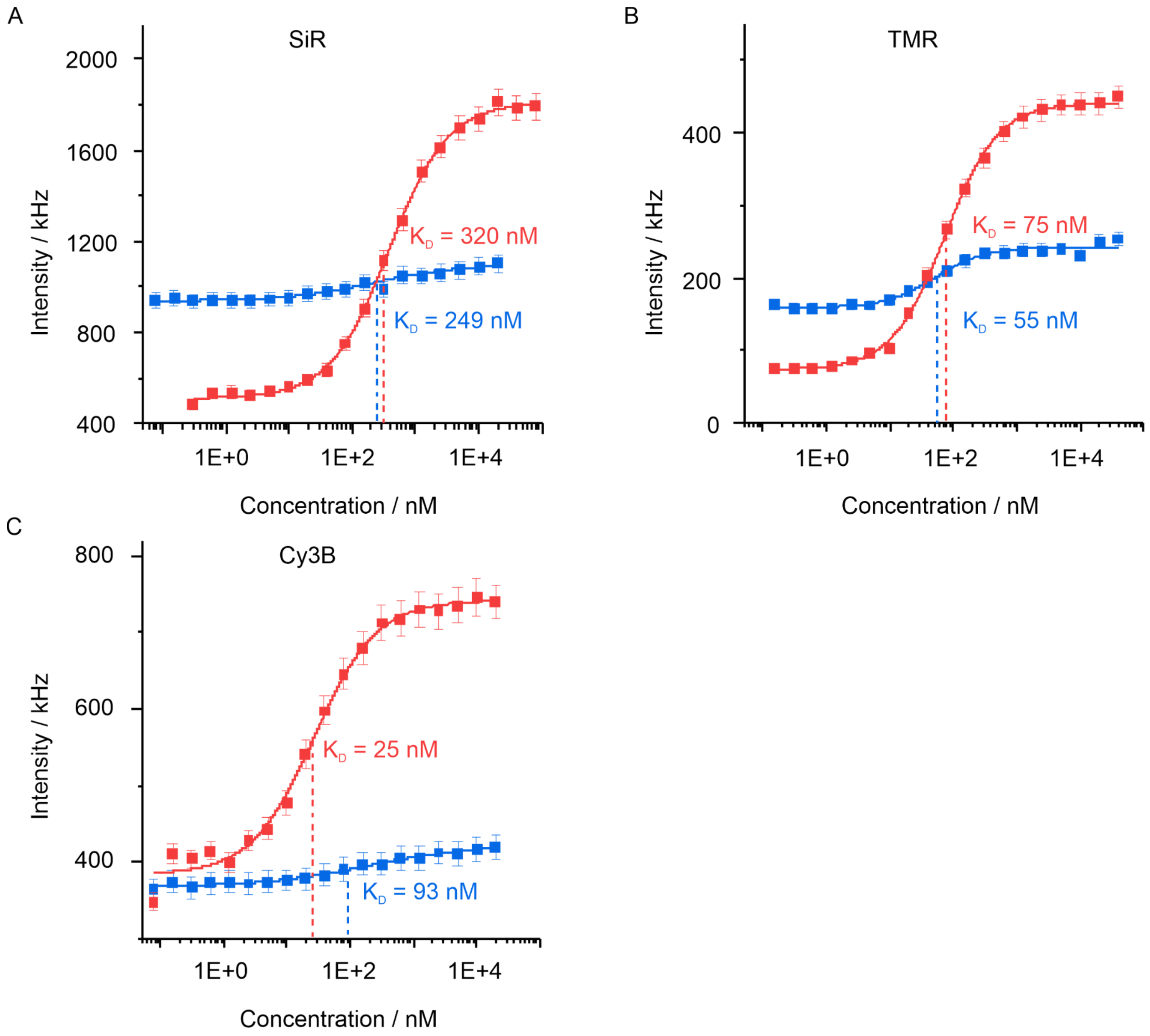


**Figure S4**: Dissociation constant determination of single- and dual-labeled imager strands with different fluorophores using fluorescent titration series. **A**. Dissociation constant measurement of P1-SiR (blue) and SiR-P1-SiR (red). **B**. Dissociation constant measurement of P1-TMR (blue) and TMR-P1-TMR (red). **C**. Dissociation constant measurement of P1-Cy3B (blue) and Cy3B-P1-Cy3B (red). Errors in the graphs represent standard deviation between three different measurements.

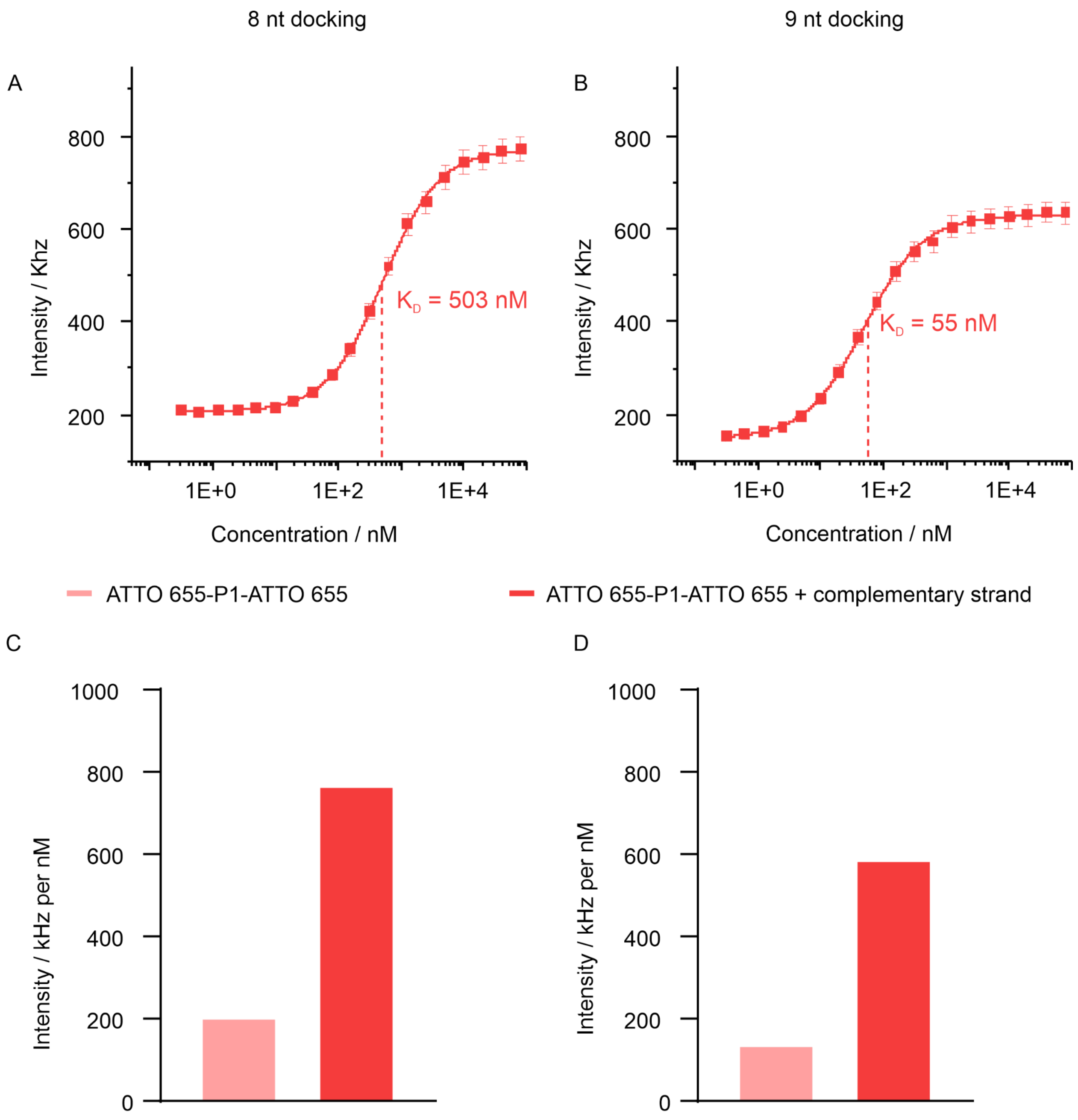


**Figure S5**: Binding affinity and fluorogenic behavior of ATTO 655-P1-ATTO 655 when hybridized to docking strands of different lengths measured using fluorescence titration series. **A**. Dissociation constant measurement of ATTO 655-P1-ATTO 655 with 8 nt long docking strands. **B**. Dissociation constant measurement of ATTO 655-P1-ATTO 655 with 9 nt long docking strands. Errors in the graphs represent standard deviation between three different measurements. **C**. Unit fluorescence intensity of single-stranded ATTO 655-P1-ATTO 655 (light red) and ATTO 655-P1-ATTO 655 hybridized to 8 nt long docking strands (red). **D**. Unit fluorescence intensity of single-stranded ATTO 655-P1-ATTO 655 (light red) and ATTO 655-P1-ATTO 655 hybridized to 9 nt long docking strands (red). Data shown in B and D were taken from Kessler et al [1].

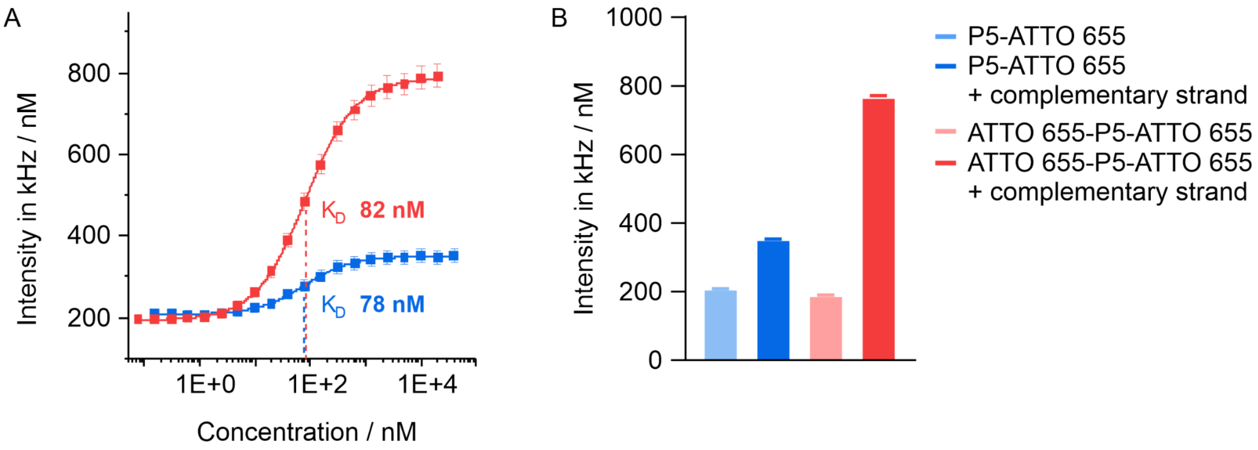


**Figure S6**: Sequence dependence on binding affinity and fluorogenic behavior measured using fluorescence titration series. **A**. Dissociation constant measurement of P5-ATTO 655 and ATTO 655-P5-ATTO 655. Errors in the graphs represent standard deviation between three different measurements. **B**. Unit fluorescence intensity of single-stranded P5-ATTO 655, ATTO 655-P5-ATTO 655 and P5-ATTO 655, ATTO 655-P5-ATTO 655 hybridized to 9 nt complementary strands.

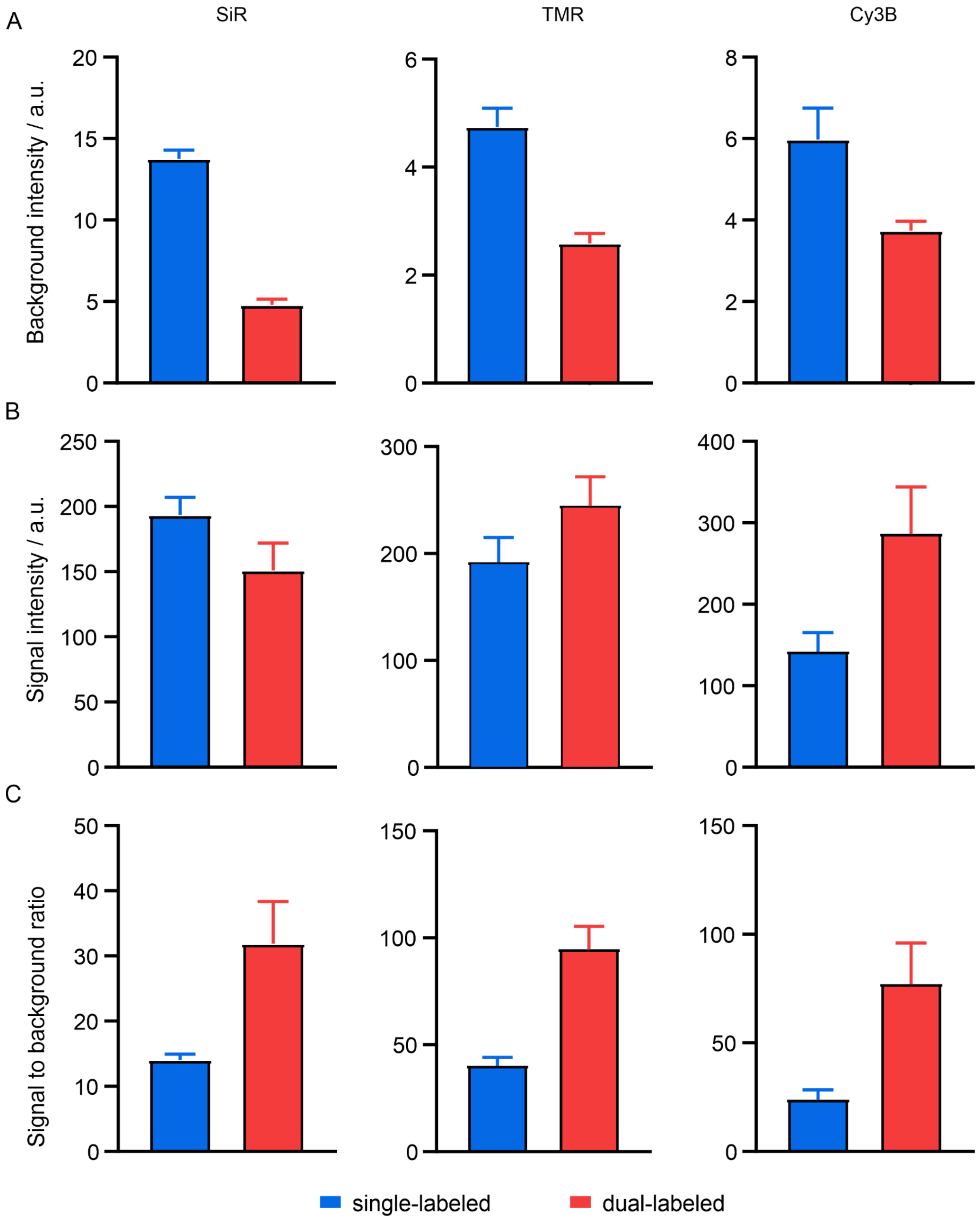


**Figure S7**: Quantification of STED imaging with single- and dual-labeled P1 imager strands with different fluorophores. Background intensity (**A**), Signal intensity (**B**) and Signal to background ratio (**C**) of single- (blue) and dual- (red) labeled P1-imager strands with SiR, TMR and Cy3B. Errors in the graphs represent standard deviation.

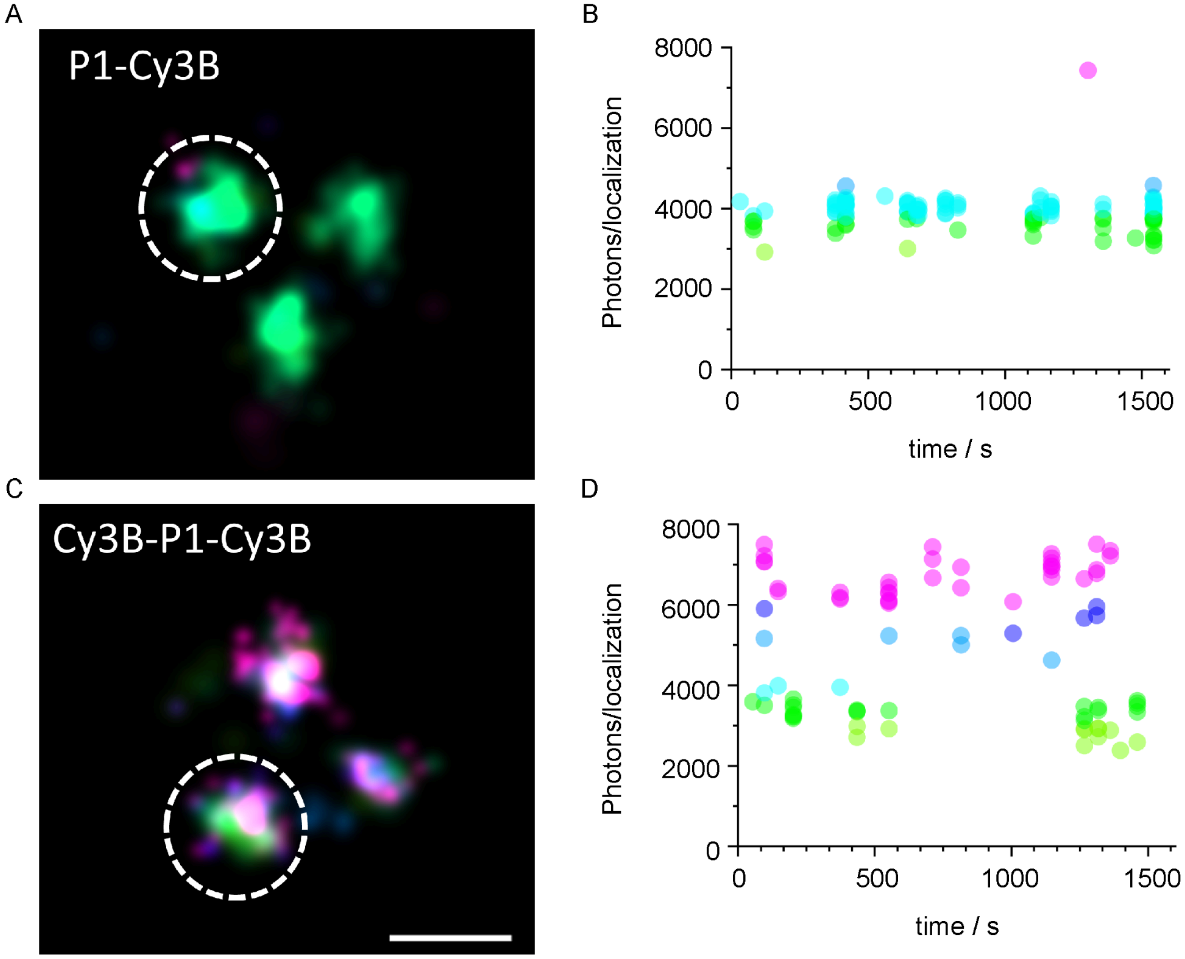


**Figure S8**: Time trace of photon emission events from measurements on DNA nanostructures. **A**. Zoom-in of one DNA-origami nanostructure showcasing the three binding sites spaced ~ 50 nm from each other imaged using DNA-PAINT with 10 nM P1-Cy3B. **B**. Photon traces over the time of the whole measurement in A. **C**. Zoom-in of one DNA-origami nanostructure showcasing the three binding sites spaced ~ 50 nm from each other imaged using DNA-PAINT with 10 nM P1-Cy3B. **D**. Photon traces over the time of the whole measurement in C. Scale bar is 50 nm.

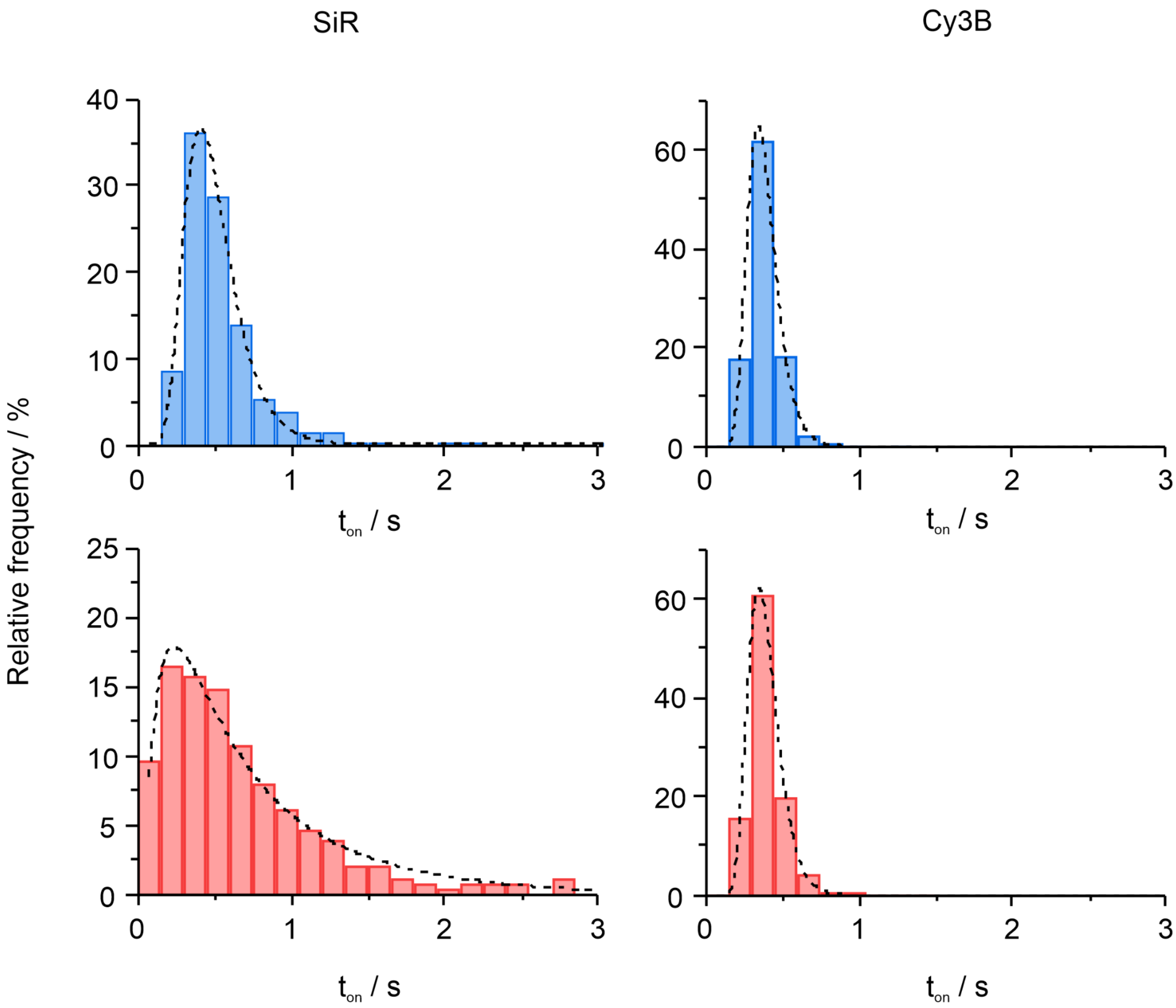


**Figure S9**: Bright times ($t_{on}$) of single- and dual-labeled imager strands with different fluorophores measured using DNA-PAINT on DNA nanostructures. $t_{on}$ of P1-SiR, P1-Cy3B and SiR-P1-SiR, Cy3B-P1-Cy3B measured using DNA-PAINT on DNA-origami nanostructures. All data were fitted using log-normal fit (**Methods**).

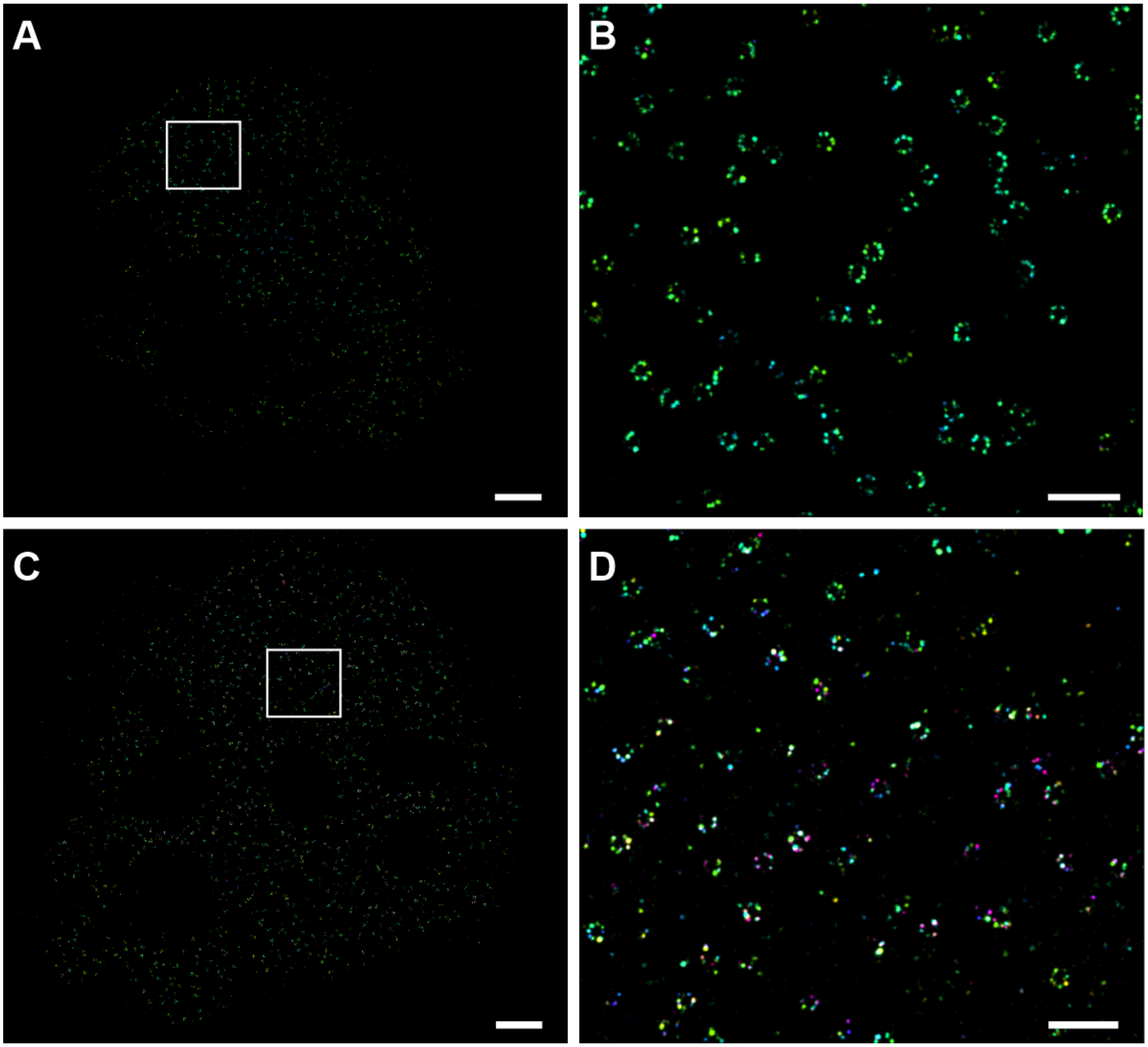


**Figure S10**: DNA-PAINT of NPCs using P1-Cy3B and Cy3B-P1-Cy3B. Whole cell overview (**A**) and zoom-in (**B**) of NPCs imaged with 1.5 nM P1-Cy3B using DNA-PAINT. Whole cell overview (**C**) and zoom-in (**D**) of NPCs imaged with 1.5 nM Cy3B-P1-Cy3B using DNA-PAINT. Scale bars for A and C are 2 µm and scale bars for B and D are 500 nm.
